# Zika virus NS4A hijacks host ANKLE2 to promote viral replication

**DOI:** 10.1101/2022.03.15.484510

**Authors:** Adam T Fishburn, Matthew W Kenaston, Nicholas J Lopez, Vivian Hoang, Traci N Shiu, Sophia T Haggard Arcé, Shahabal S Khan, Priya S Shah

**Affiliations:** University of California, Davis, Department of Microbiology and Molecular Genetics; University of California, Davis, Department of Chemical Engineering

**Keywords:** Zika virus, flavivirus, NS4A, ANKLE2, virus-host interaction, microcephaly

## Abstract

Zika virus (ZIKV) is infamous among flaviviruses for its unique association with congenital birth defects, notably microcephaly. We previously mapped ZIKV-host protein interactions and identified the interaction between ZIKV NS4A and host ANKLE2, which itself has established ties to congenital microcephaly. In fruit flies, NS4A induces microcephaly phenotypes in an ANKLE2-dependent manner. This suggests that NS4A interacts with ANKLE2 to dysregulate cell behavior and contributes to abnormal host neurodevelopment. Here, we explore the role of ANKLE2 in ZIKV replication to understand the biological significance of the interaction from the viral perspective. We show that knockdown of ANKLE2 reduces replication of two ZIKV strains, across multiple MOIs and timepoints. We observe that localization of ANKLE2 is drastically shifted to sites of NS4A accumulation during infection. We investigate which domains of ANKLE2 mediate this behavior and the interaction with NS4A. Using co-immunoprecipitation, we show that deletion of either the transmembrane or LEM domain has little impact on the interaction, but deletion of both significantly reduces interaction with NS4A. We show that the C-terminal transmembrane domains of NS4A stabilize the interaction with ANKLE2. Finally, we explore this interaction in other flaviviruses and observe ANKLE2 interacts with NS4A across four additional mosquito-borne flaviviruses. Together, these results suggest NS4A interacts with ANKLE2 through a combination of its transmembrane and LEM domains, bringing it to sites of ZIKV replication to promote replication through an unknown mechanism. Taken together with our previous results, our findings indicate that, in the process of hijacking ANKLE2 for replication, ZIKV disrupts its physiological function to cause disease.

**Importance:** The ZIKV epidemic led to the astonishing revelation that congenital ZIKV infection is associated with devastating birth defects, including microcephaly. Microcephaly is the condition in which head and brain size are severely reduced, and is often accompanied by intellectual disability. The molecular mechanisms by which ZIKV replicates and causes microcephaly are still incompletely understood. We previously identified the protein interaction between ZIKV NS4A and host ANKLE2, which is associated with congenital microcephaly. In flies, NS4A induces microcephaly in an ANKLE2-dependent manner, suggesting this interaction is crucial for ZIKV pathogenesis. Here, we explore the relevance of this physical interaction for virus replication. We find that ANKLE2 promotes ZIKV replication, concentrates at sites of NS4A accumulation during infection, and interacts with NS4A via its N-terminal domain. Thus, this represents a rare example of a ZIKV-host protein interaction that impacts both disease and virus replication.

## Introduction

Zika virus (ZIKV) is a positive sense RNA virus belonging to the genus *Flavivirus* and can cause severe disease. ZIKV infection in otherwise healthy adults typically leads to mild symptoms and very rarely Guillain-Barré Syndrome (1). The primary concern surrounding ZIKV arises from the occurrence of Congenital Zika Syndrome (CZS) in individuals born from mothers infected during pregnancy (2). CZS is a spectrum of disease and can be clinically characterized by multiple hallmark features, including congenital contractures, ocular anomalies, cortical calcifications, and in the most severe cases, microcephaly (3, 4).

Microcephaly is a condition in which the head and brain size are significantly reduced at birth and is associated with a wide range of complications, including developmental delays, intellectual disability, and predisposition to seizures (5). Beyond CZS, microcephaly can arise from genetic mutations (6), exposure to toxins during pregnancy (7–9), and congenital infections by pathogens such as rubella virus (10, 11), cytomegalovirus (12), and *Toxoplasma gondii* (13).

The specific molecular mechanisms by which ZIKV contributes to abnormal neurodevelopment and microcephaly are not entirely understood. ZIKV can infect placental tissue and compromise placental function (14–16), potentially providing ZIKV access to the fetal compartment. Gestational timing and type I interferon responses *in utero* are also critical in ZIKV susceptibility (17, 18). Upon fetal infection, ZIKV also breaches the blood-brain barrier and infects susceptible neuronal tissue (19–23). Infection of these neuronal tissues is associated with direct tissue damage through the cytopathic effect (24–26), molecular level pathogenesis (27), disruption of cell division (24, 28–30), and dysregulation of developmental pathways (31–37).

Previously, we used affinity-purification and mass-spectrometry to identify ZIKV-host protein-protein interactions that may contribute to the development of microcephaly in CZS (38). By searching for host proteins with known roles in neurodevelopment or associations with microcephaly, we identified the interaction between the ZIKV non-structural protein 4A (NS4A) and host Ankyrin-repeat and LEM domain containing protein 2 (ANKLE2). ANKLE2 has well established roles in human congenital microcephaly, which has been replicated *in vivo* using *Drosophila* (39–41). Mutations in *ankle2* result in small brain phenotypes and cellular defects in neuroblasts in 3^rd^ instar larvae. These phenotypes are rescued by the expression of human ANKLE2, suggesting that human ANKLE2 and *Drosophila* Ankle2 are functionally conserved in brain development (39, 40). Using this *Drosophila* model, we showed that transgenic expression of ZIKV NS4A induces similar microcephaly phenotypes which are also rescued by the expression of human ANKLE2. Overall, this suggests NS4A induces microcephaly *in vivo* in an ANKLE2-dependent manner (38, 40).

Flaviviruses physically hijack host factors for their own replication and use many virus-host protein interactions to reshape the host cell into a niche that maximizes virus replication (42). Whether NS4A inhibition of ANKLE2 function during development is simply an unfortunate coincidence, or if there is a functional role for this virus-host protein interaction in ZIKV replication is unknown. ANKLE2 is primarily considered a scaffolding protein, facilitating protein-protein interactions between kinases, phosphatases, and their substrates. ANKLE2 localizes to the ER and inner nuclear membrane where it mediates interactions with proteins, including barrier to autointegration factor (BANF1), vaccinia related kinase 1 (VRK1), and protein phosphatase 2A (PP2A), to control nuclear membrane disassembly during cell division (43, 44). NS4A itself is integral to the formation of flavivirus replication compartments in the ER (45, 46) and facilitates other aspects of flavivirus replication through interactions with host proteins (47–50). We hypothesize that NS4A interacts with ANKLE2 to hijack its scaffolding function in a way beneficial to virus replication.

In this study we explore the role of ANKLE2 in ZIKV replication and what structural domains mediate the biophysical interaction between ANKLE2 and NS4A. We show that knockdown of ANKLE2 reduces ZIKV replication across multiple ZIKV strains and MOIs. We find ANKLE2 is concentrated at sites of NS4A accumulation during infection. We identify that the transmembrane (TM) domain of ANKLE2 localizes it to the ER and nuclear lamina, but both the TM and LEM domains are necessary for its physical interaction with NS4A. Finally, we establish the domain of NS4A that interacts with ANKLE2 and show the conservation of the ANKLE2-NS4A interaction and this domain among other mosquito-borne flaviviruses. Together, we report the novel function of ANKLE2 in promoting ZIKV infection, providing evidence that NS4A disruption of neurodevelopment through ANKLE2 may arise from an underlying virus replication mechanism.

## Results

### Knockdown of ANKLE2 reduces ZIKV replication

To explore the role of ANKLE2 in ZIKV replication, we used a gene perturbation approach. We depleted ANKLE2 using CRISPRi with two synthetic guide RNAs (gRNAs) (Supplementary Figure 1A). Huh7 cells stably expressing dCas9 were generated using lentiviral transduction (Supplementary Figure 1B). We chose Huh7 cells, derived from a hepatocarcinoma, since they are readily infected by ZIKV *in vitro* (51, 52), and the liver is an established site of ZIKV replication *in vivo* (53–55). CRISPRi gRNAs were delivered by reverse transfection. At 72 hours post transfection (hpt) the expression of ANKLE2 in ANKLE2 knockdown cells was decreased to ∼30% of negative control RNA (ncRNA) (Figure 1A and B). *ANKLE2* mRNA was also significantly reduced by both gRNAs compared to ncRNA (Supplementary Figure 1C). We confirmed that cell viability of ANKLE2 knockdown cells was comparable to that of cells transfected with ncRNA (Supplementary Figure 1D).

**Figure 1:**
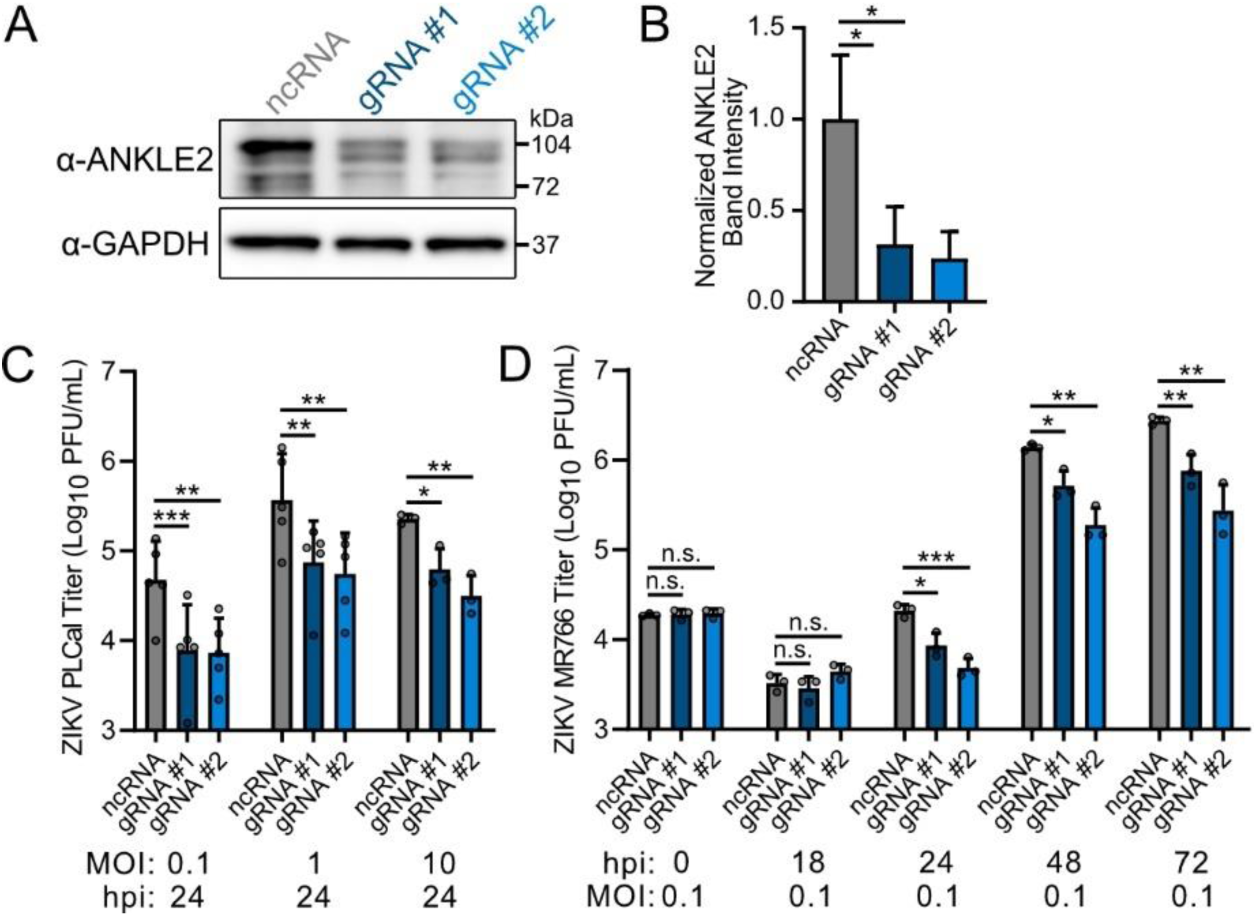
Knockdown of *ANKLE2* reduces ZIKV replication across multiple MOIs. **A)** Western blot of ANKLE2 following CRISPRi knockdown (72 hpt). **B)** Densitometry measurement of ANKLE2 band intensity, normalized to GAPDH. Data shown from three biological replciates. **C)** ZIKV PLCal titers at 24 hpi across three MOIs. **D)** ZIKV MR766 titers at MOI 0.1 over 72 hpi. All error bars represent standard deviation. Assessment of ZIKV PLCal at MOI 0.1 and 1 were done with five paired biological replicates. The remaining conditions were done with three technical replicates. Student’s two-tailed T-test, paired for PLCal MOI 0.1 and 1, unpaired for remaining conditions. * p < 0.05, ** p < 0.01, *** p < 0.001.

To determine if ANKLE2 plays a role in ZIKV replication, we infected Huh7-dCas9 cells with ZIKV after 72 hours of ANKLE2 knockdown. Infection with ZIKV PLCal (Thailand, 2013) revealed modest (maximum 7-fold decrease) but consistent and statistically significant decreases in virus titers between ncRNA and *ANKLE2* gRNA treated conditions at 24 hours post infection (hpi) for a range of multiplicities of infection (MOIs) (Figure 1B). To further support the role of ANKLE2 in ZIKV replication we repeated this experiment using the ZIKV strain MR766 (Uganda, 1947) and this time measured virus titers at multiple timepoint post-infection for a single MOI. Western blot analysis of uninfected cells showed that ANKLE2 knockdown persists for 6 days post transfection, which would span the entire experimental time course for this experiment (Supplementary Figure 1E). Following knockdown and infection, we observed consistent decreases in virus titers starting at 24 hpi, with a maximum 10-fold decrease at 72 hpi (Figure 1D). Together, this shows knockdown of *ANKLE2* reduces ZIKV replication in Huh7 cells.

### ANKLE2 colocalizes with NS4A and dsRNA during infection

ANKLE2 is known to localize to the ER and inner nuclear membrane where it facilitates host protein-protein interactions (43). NS4A is essential for remodeling the host ER into a niche suited for flavivirus replication. NS4A alone can induce membrane curvature which promotes the formation of virus replication compartments (46). It can also hijack host proteins to further facilitate ER remodeling (50). Given our previous proteomics that established the interaction between ANKLE2 and NS4A (38) we speculated that the scaffolding function of ANKLE2 may be hijacked by NS4A to facilitate replication. We therefore evaluated ANKLE2 subcellular localization during ZIKV infection. Due to a lack of compatible immunofluorescence-grade antibodies we generated HEK293T cell lines that express FLAG affinity tagged fusions of ANKLE2 (ANKLE2-FLAG). As controls, we also generated cell lines that express GFP-FLAG as a general non-specific control and ANKLE1-FLAG to distinguish between intrinsic localization in this family of proteins, and localization unique to ANKLE2. We infected these cells with several ZIKV strains (PLCal, MR766, and PRVABC59) and evaluated the localization of FLAG and NS4A. Strikingly, we found that ANKLE2 distribution was concentrated to clusters that had near perfect co-localization with clusters of NS4A during infection (Figure 2A). Conversely, we found that the localization and behavior of GFP and ANKLE1 did not change following ZIKV infection (Figure 2B-C). We measured colocalization between FLAG and NS4A signal using Pearson’s correlation and consistently found very high levels of colocalization between ANKLE2 and NS4A (Pearson’s R-value = 0.93 ± 0.04 across all 3 strains), which was significantly higher than GFP or ANKLE1 with NS4A (R-value = 0.43 ± 0.10, 0.46 ± 0.15, respectively) (Figure 2D).

**Figure 2:**
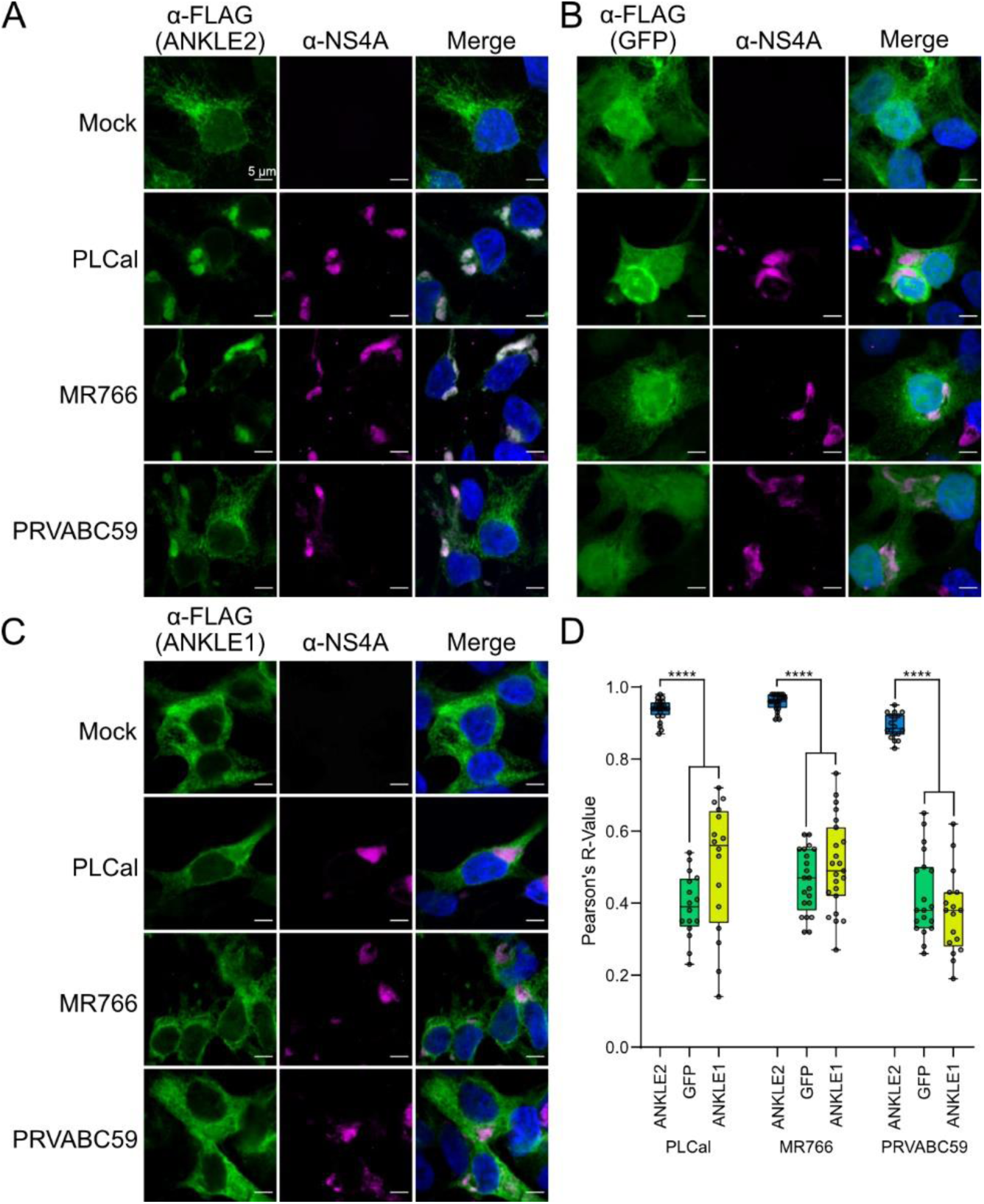
ANKLE2 localization pattern is altered during ZIKV infection and colocalizes with NS4A. **A-C)** HEK293T cells expressing ANKLE2, GFP, or ANKLE1 were infected with noted ZIKV strain for 48 hours. Proteins were visualized by immunofluorescence and confocal microscopy: FLAG (green) and NS4A (magenta) and Hoechst (blue). All scale bars = 5 µm. **D)** Pearson’s correlation was measured between the FLAG and NS4A signal in infected cells. Five images and n = 16-24 cells measured per condition. One-way ANOVA with Šidák multiple comparisons test, **** p < 0.0001.

Given the colocalization of ANKLE2 and NS4A during infection, the positive role of ANKLE2 in the ZIKV replication cycle, and the known role of NS4A in forming ZIKV replication factories, we next explored if ANKLE2 colocalizes with dsRNA generated in replication factories during viral genome replication. We and others observed that ZIKV MR766 had the highest replicative efficiency in HEK293T cells and therefore used it for our remaining immunofluorescence experiments (56). To evaluate the colocalization of ANKLE2 with sites of ZIKV replication we used antibodies against endogenous ANKLE2 and flavivirus envelope (E) protein (4G2) or dsRNA (rJ2) (Figure 3A-B). We observed high colocalization between ANKLE2 and E (R-value = 0.89 ± 0.05) that clustered similarly in cells to FLAG-tagged ANKLE2 when infected with the same ZIKV MR766 strain (Figure 3G). We observed that clusters of ANKLE2 signal in ZIKV infected cells partially overlapped with sites of dsRNA signal, although quantitative measurement revealed only modest colocalization (R-value = 0.44 ± 0.10), again, likely due to the punctate behavior of dsRNA. To confirm these were indeed sites of ZIKV genome replication we performed the same experiment evaluating the colocalization between NS4A and E or dsRNA, since NS4A is inherently involved in forming replication factories where dsRNA is formed (Figure 3C-D). This revealed very high correlation between NS4A and E (R-value = 0.96 ± 0.02), suggesting these large clusters maybe the location where viral protein accumulates during or after translation. Colocalization between NS4A and dsRNA was lower (R-value = 0.42 ± 0.12) and similar to that of ANKLE2 with dsRNA. In fact, there was no statistically significant difference between them (Figure 3H). As an additional control we compared E and dsRNA colocalization with GFP. GFP had a substantially and significantly lower colocalization than ANKLE2 or NS4A with both E and dsRNA (Figure 3E-H), suggesting that the modest correlation observed for ANKLE2 and NS4A with dsRNA is biologically meaningful. We performed the same analysis in Huh7 cells and observed similar clustering by ANKLE2 in infected cells, although quantitative measurements of colocalization were lower in these cells (Supplementary Figure 2). Collectively, these results suggest that during ZIKV infection, ANKLE2 is present at sites of viral dsRNA formation.

**Figure 3:**
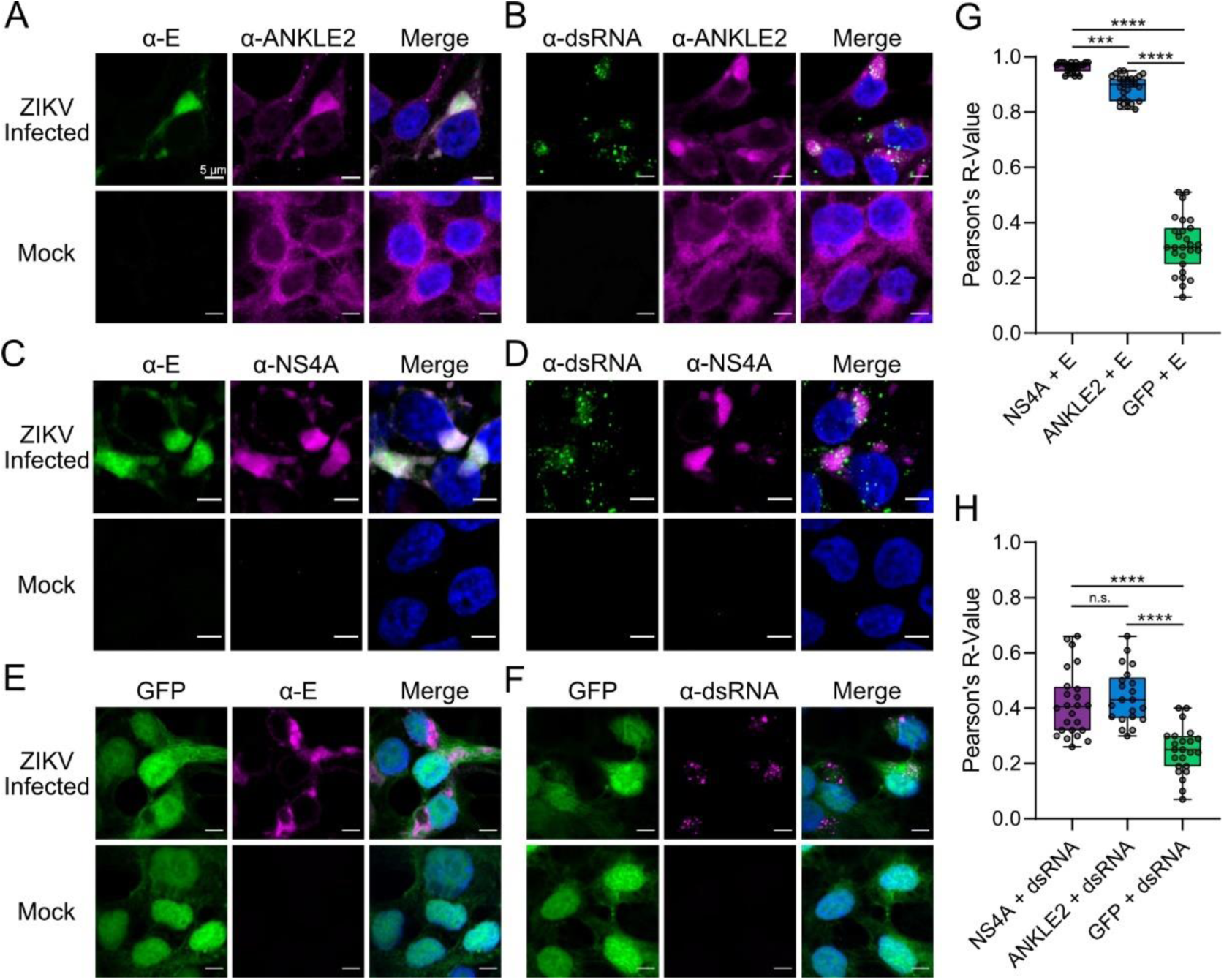
ANKLE2 colocalizes with ZIKV proteins and dsRNA. **A-F)** HEK293T cells were infected with ZIKV MR766 (MOI 5) for 48 hours prior to fixation and antibody staining. All scale bars = 5 µm. **G-H)** Pearson’s R-value measured using ImageJ colocalization tool of n = 21-27 cells. One-way ANOVA with Tukey’s multiple comparisons test, n.s. = not significant, *** p < 0.001, **** p < 0.0001.

### NS4A interacts with ANKLE2 through its TM and LEM domains

To further understand the biophysical interaction between NS4A and ANKLE2, we sought to determine which domains of ANKLE2 were necessary for the interaction. The structure of ANKLE2 has not yet been resolved. Thus, we employed the structural prediction provided by AlphaFold2 (57, 58). This revealed high-confidence structured regions corresponding to the known TM, LEM, and ankyrin repeat (ANK) domains. Surprisingly, this also revealed three previously uncharacterized high-confidence structures (Supplementary Figure 3). Using this predicted structure as a template, we then generated seven C-terminal truncation mutants with progressively fewer of these domains or structures, each with FLAG affinity tags (3xF) (Supplementary Figure 4A). To characterize the biochemical behavior of these truncations we performed subcellular fractionation to enrich for cytosolic, membrane, and nuclear/nuclear lamina compartments. Western blotting analysis of these fractions showed the presence of each primarily in the membrane-bound (including ER, Golgi, etc.) and nuclear/nuclear-lamina fractions (Supplementary Figure 4B and C). This result is expected given the established localization of ANKLE2 to the ER and inner nuclear membrane (43, 44), and the role of the TM, which was not deleted, in facilitating ER localization (59). To evaluate interaction of these ANKLE2 truncations with NS4A we co-transfected each with C-terminally Strep-tagged NS4A-2K plasmid we generated previously (38) and then performed FLAG co-immunoprecipitation (co-IP). Intriguingly, FLAG co-IP revealed that each truncation maintained its interaction with NS4A-2K, although Δ54-938 only very weakly interacted (Supplementary Figure 4D). Together, these results showed the C-terminal deletions of ANKLE2 do not substantially impact its localization or interaction with ZIKV NS4A.

Given that C-terminal deletions did not impact ANKLE2 or its interaction with NS4A, we generated additional mutants with either N-terminal deletions of the TM domain only (Δ2-53), the TM and LEM domains (Δ2-157), the LEM domain only (Δ54-158), or the LEM through ANK domains (Δ54-474) (Figure 4A). We performed similar subcellular fractionation analysis which showed that Δ2-53 and Δ2-157 were enriched in the cytosolic fraction and depleted in the nuclear/nuclear lamina fraction, but maintained some membrane (ER) localization (Figure 4B-C). Confocal microscopy of Δ2-53 and Δ2-157 showed disperse signal in the cytosol with decreased overlap with the ER marker Calreticulin, whereas WT ANKLE2 and internal deletions Δ54-158 and Δ54-474 had high colocalization with the ER as measured by Pearson’s correlation (Figure 4D-E). Thus, both ER and nuclear/nuclear lamina localization were disrupted in the Δ2-53 and Δ2-157 mutant.

**Figure 4:**
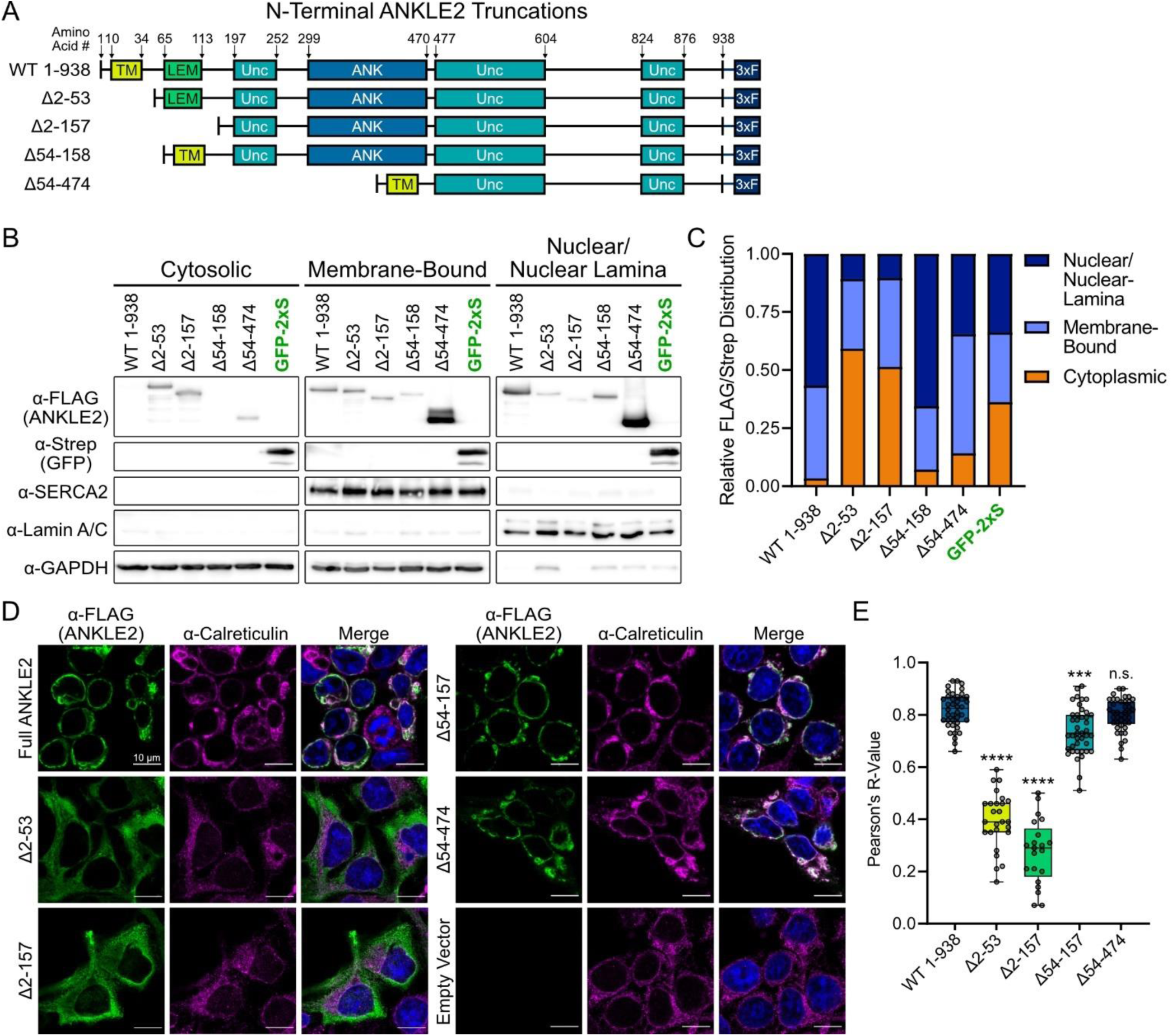
Assessment of ANKLE2 N-terminal truncations and internal deletion mutants. **A)** N-terminal ANKLE2 truncations and internal deletion mutants were designed based on a structural prediction by AlphaFold2. TM = transmembrane domain; LEM = LAP2, emerin, MAN1 domain; Unc = uncharacterized structure predicted by AlphaFold2; ANK = ankyrin repeat domain; 3xF = 3x FLAG affinity tag. **B)** Western blot analysis of subcellular fractions of cells transfected with the indicated ANKLE2 construct. SERCA2 and Lamin A/C antibodies were used to validate enrichment of membrane-bound and nuclear/nuclear lamina fractions, respectively, while GAPDH was used as a cytoplasmic marker. **C)** Densitometry analysis was performed on western blot images by measuring the intensity of FLAG/Strep bands relative to each fraction’s marker. These values were then plotted as a fractional distribution for relative intensity across all fractions. **D)** Confocal microscopy of HEK293T cell transfected with noted ANKLE2-3xFLAG truncations. The colocalization of each truncation (green) was compared with the ER marker, Calreticulin (magenta). **E)** Pearson’s correlation was quantified from n = 21-41 cells across 5-11 images per condition. All scale bars = 10 µm. One-way ANOVA with Dunnett’s multiple comparisons test compared to control (WT ANKLE2 1-938), n.s. = not significant, *** p < 0.001, **** p < 0.0001.

Co-transfection with NS4A-2K and FLAG co-IP of these ANKLE2 mutants revealed that deletion of both the TM and LEM domains (Δ2-157) strongly ablated interaction with NS4A-2K, whereas deletion of either the TM or LEM domain alone retained the interaction with NS4A-2K (Figure 5A, Supplementary Figure 4D). This was somewhat surprising since deletion of the TM substantially altered subcellular localization that could preclude the interaction with NS4A in cells. To corroborate this biochemical finding, we visualized the subcellular localization of these ANKLE2 truncations along with NS4A during infection. Here, we clearly observed similar clusters of NS4A that colocalized with wild-type ANKLE2, but not the Δ2-53 and Δ2-157 mutants. Instead, we observed our Δ2-53 and Δ2-157 ANKLE2 mutants appeared to be distinctly separated from sites of NS4A accumulation (Figure 5B-C, white arrows). However, this separation is not necessarily indicative of the ability to physically interact, as Δ2-53 still readily interacts with NS4A when assayed by co-IP since subcellular compartments can be mixed during cell lysis. Altogether, these results suggest that while the TM domain controls ER and nuclear lamina localization of ANKLE2, both the TM and LEM regions together contribute to the interaction with NS4A, with presence of at least one being sufficient for the biochemical interaction.

**Figure 5:**
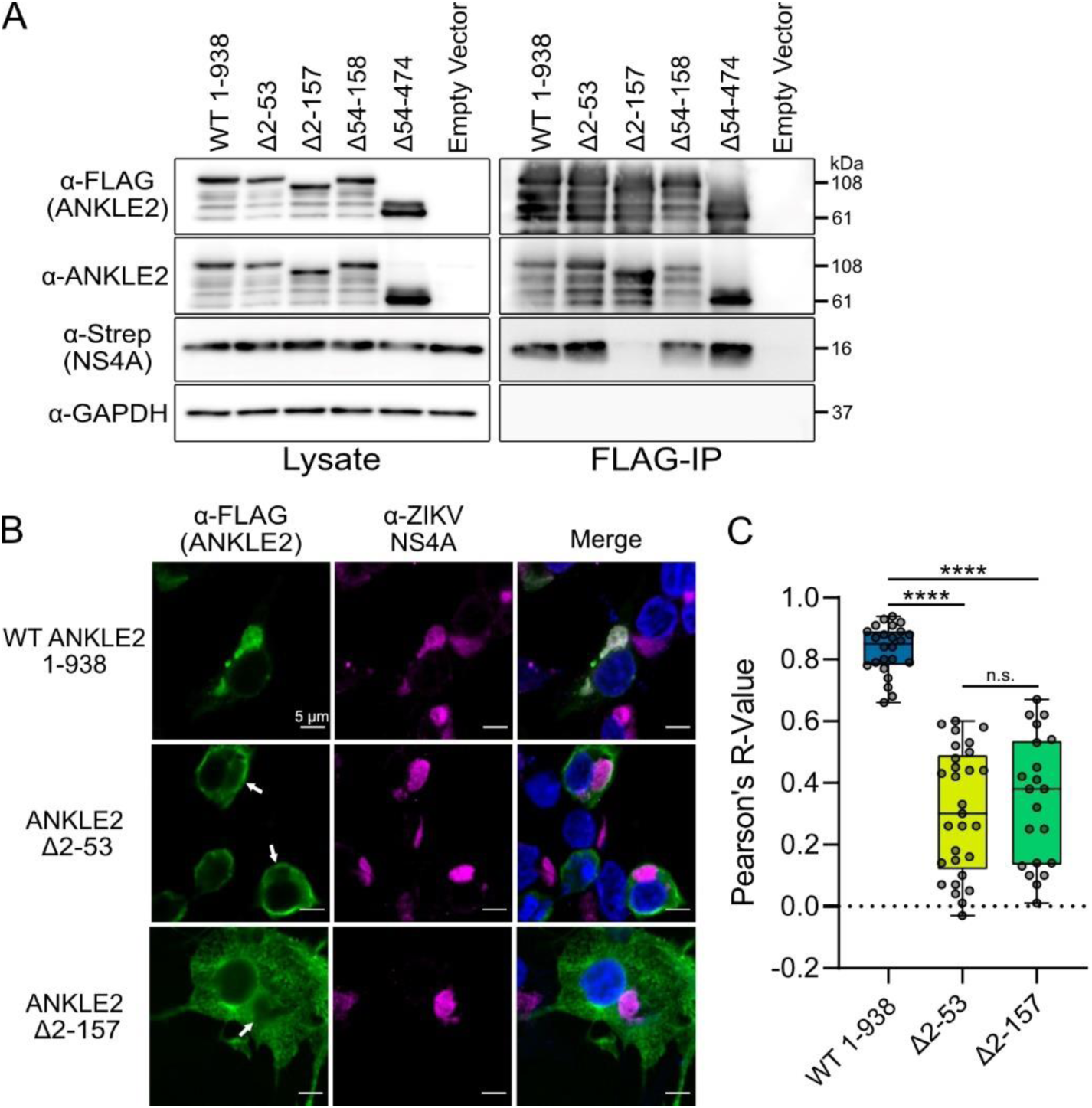
ANKLE2 truncations reveal that ER localization, and the LEM domain are important for interaction with NS4A-2K. **A)** Western blotting of lysate and FLAG-IP samples. Empty vector condition denotes transfection with NS4A-2K-2xStrep plasmid and an empty plasmid instead of one harboring an ANKLE2-3xF sequence to account for potential FLAG-IP background. ANKLE2-3xFLAG truncations were detected with both FLAG and ANKLE2 antibodies to ensure that lower molecular weight bands are degradation products of these truncations. Due to high transfection efficiency of our truncations, endogenous ANKLE2 is only observable with higher exposure and is only faintly visible. **B)** HEK293T cells expressing indicated ANKLE2 construct and infected with ZIKV were analyzed by immunofluorescence microscopy for ANKLE2-3xFLAG (green) and NS4A (magenta) localization, and Hoescht nuclear stain (blue). White arrows denote regions where truncated ANKLE2 is absent from clusters of NS4A. **C)** Pearson’s correlation was quantified from n = 21-29 cells across 5 images per condition. All scale bars = 5 µm. One-way ANOVA with Tukey’s multiple comparisons test, n.s. = not significant, **** p < 0.0001.

### NS4A-ANKLE2 interaction is dependent on NS4A TM2 and TM3

To further explore the determinants of the NS4A-ANKLE2 interaction, we sought to generate NS4A truncation mutants. Using a previous evaluation of dengue virus (DENV) NS4A as a template (46), we initially generated plasmids encoding one of three C-terminal truncations, each removing an additional TM domain (Figure 6A). As done previously with our ANKLE2 truncations, we characterized the localization of these NS4A truncations using subcellular fractionation and confocal microscopy. This confirmed that each was retained in the membrane-bound fraction (Figure 6B and C), and all had similar colocalization with the ER-marker Calnexin (Figure 6D and E). We then expressed these NS4A truncations in HEK293T cells and performed co-IP against the Strep affinity tag to determine interaction with endogenous ANKLE2 (Figure 7A). The first of these truncations, NS4A ΔTM4, had better expression and interaction with ANKLE2 than our NS4A-2K. This also represents a biologically relevant form of NS4A during ZIKV infection, since the 2K-peptide is first cleaved at the C-terminus of NS4A to separate NS4A from 2K-NS4B (60). The other truncations, NS4A ΔTM3-4 and ΔTM2-4, had significantly reduced to no visible interaction with ANKLE2, suggesting that at least TM3 is crucial for stabilizing the interaction with ANKLE2. To corroborate this result, as well as our previous results with ANKLE2 (Figure 5A), we co-transfected our ANKLE2-3xFLAG and NS4A-2xStrep truncation constructs and performed FLAG Co-IP (Figure 7B). Here, to display the importance of NS4A TM domains, we additionally generated a fourth NS4A truncation that does not express any TM domains (NS4A ΔTM1-4). In this experiment we observed that NS4A ΔTM3-4 had detectable but dramatically reduced interaction with ANKLE2. However, NS4A ΔTM2-4 had no detectable interaction. Additionally, ΔTM1-4 had very low expression and we could not draw conclusions regarding its ability to interact with ANKLE2. Our co-IP results indicate that both TM2 and TM3 contribute to the interaction with ANKLE2. Finally, we evaluated the effect of the 2K-peptide on the interaction between NS4A and ANKLE2 Δ2-158, which had strongly reduced interaction with NS4A-2K previously (Figure 5A). We observed faint and reduced interaction between ANKLE2 Δ2-157 and NS4A-2K, but no detectable interaction between ANKLE2 Δ2-157 and NS4A ΔTM4, suggesting that the 2K-peptide may stabilize the interaction outside of the TM and LEM domains, but that the processed form of NS4A requires these ANKLE2 domains for interaction.

**Figure 6:**
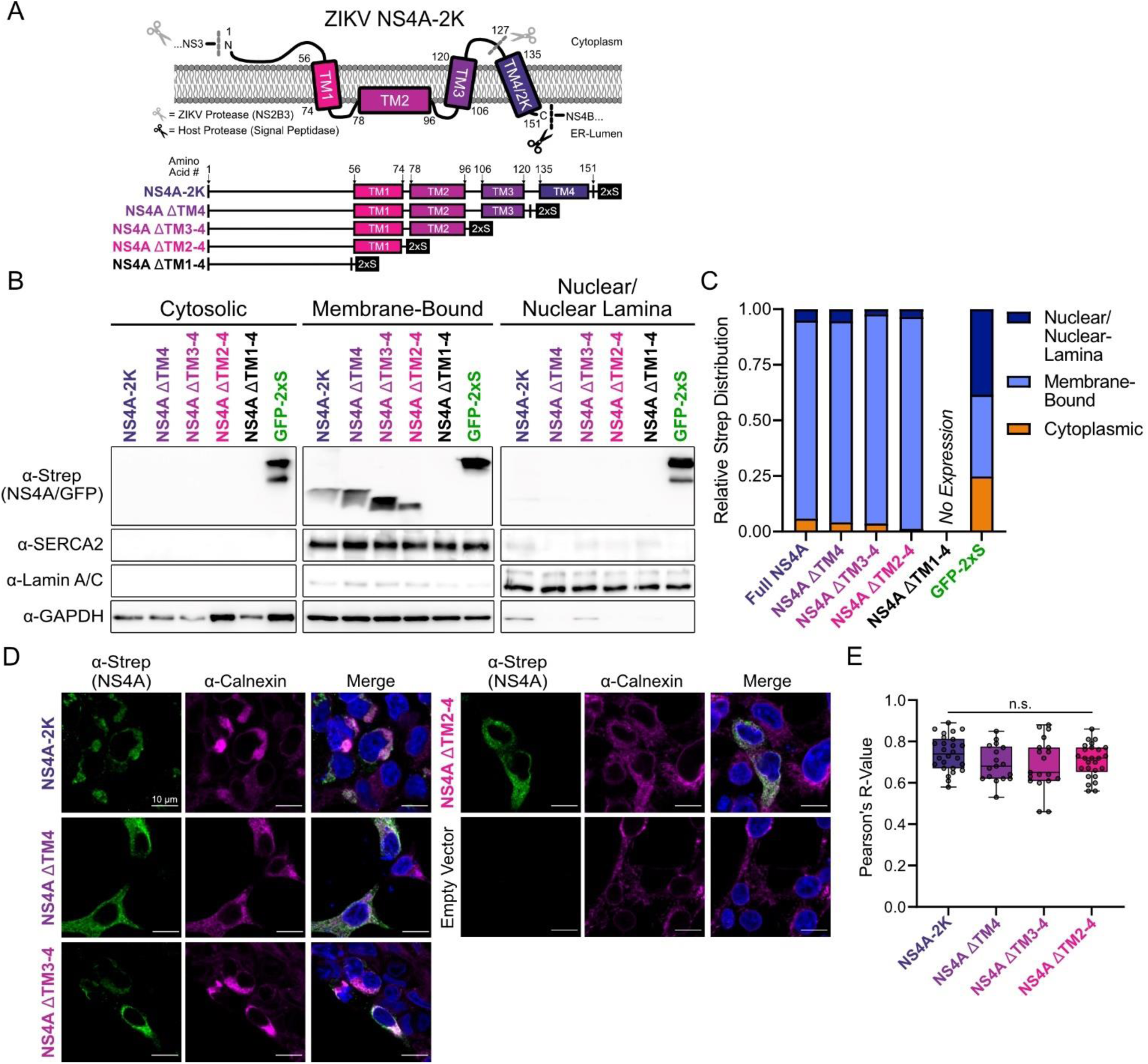
ZIKV NS4A C-terminal truncations are localized to the ER. **A)** Schematic of NS4A with annotated amino acid number for consensus predicted transmembrane domains (above). C-terminal truncations mutants were generated based on these domains, each with C-terminal 2xStrep affinity tags (below). TM = transmembrane domain, 2xS = 2xStrep affinity tag. **B)** Western blot analysis of subcellular fractions of cells transfected with the indicated NS4A construct. SERCA2 and Lamin A/C antibodies were used to validate enrichment of membrane-bound and nuclear fractions, respectively, while GAPDH was used as a cytoplasmic marker. **C)** Densitometry analysis was performed on western blot images by measuring the intensity of Strep bands relative to each fraction’s marker. These values were then plotted as a fractional distribution for relative intensity across all fractions. **D)** Confocal microscopy of HEK293T cell transfected with noted NS4A-2xS truncations. The colocalization of each truncation (green) was compared with the ER marker, calnexin (magenta). **E)** Pearson’s correlation was quantified from n = 17-26 cells across 5 images per condition. All scale bars = 10 µm. One-way ANOVA, n.s. = not significant.

**Figure 7:**
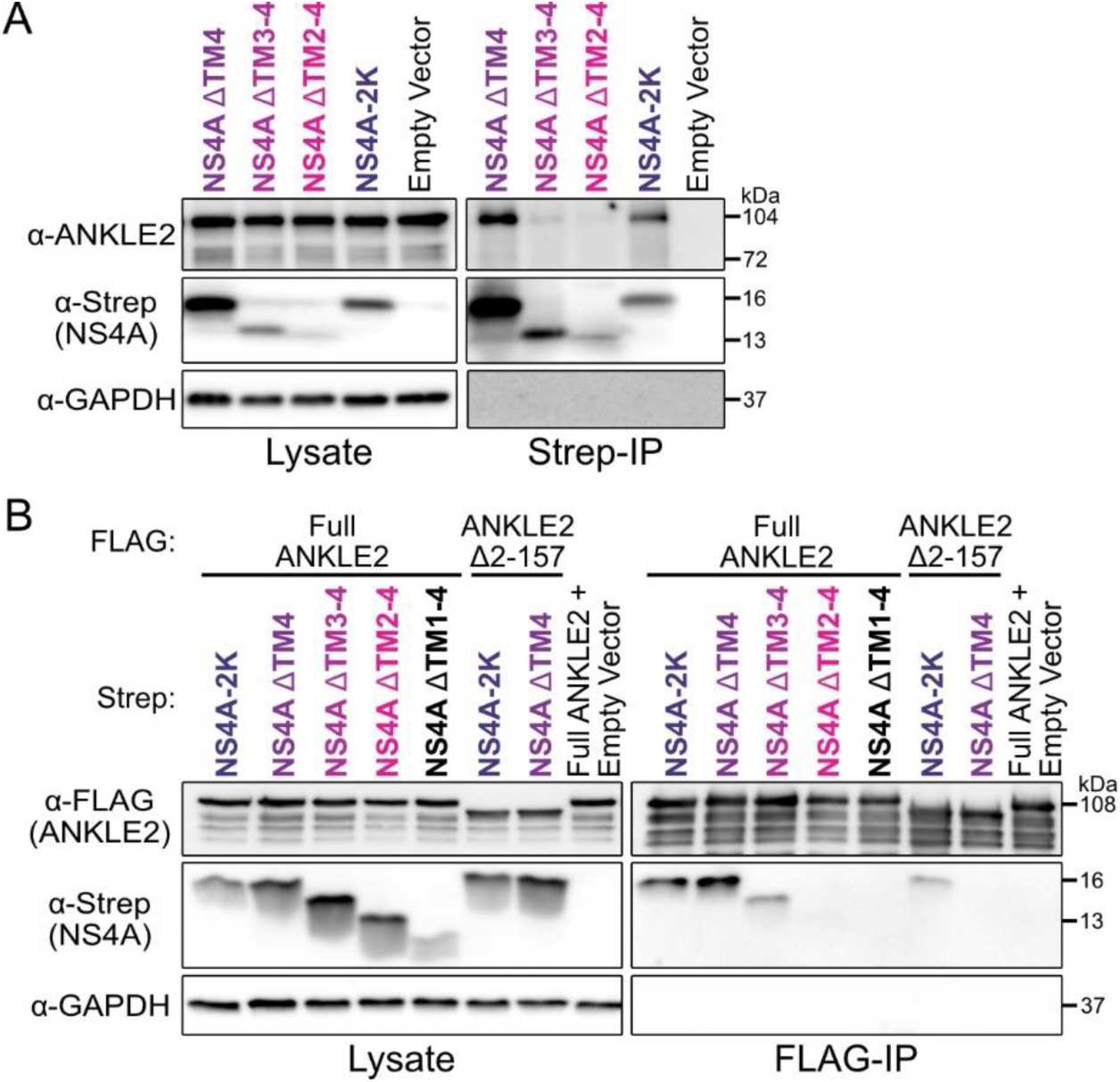
TM2 and TM3 of NS4A mediate interaction with ANKLE2 in vitro. A) Western blotting of lysate and Strep IP samples with indicated NS4A. B) Western blotting of lysate and FLAG IP samples of cells co-transfected with indicated ANKLE2 and NS4A.

### Conservation of flavivirus NS4A and its interaction with ANKLE2

Given that ANKLE2 promotes ZIKV replication, we next sought to explore the conservation of this interaction amongst other mosquito-borne flaviviruses. We generated NS4A Strep-tagged fusions for these four additional flaviviruses (dengue virus [DENV], West Nile virus [WNV], yellow fever virus [YFV], and Japanese encephalitis virus [JEV]) and evaluated their interaction with ANKLE2 using the reverse FLAG-IP we performed previously (Figure 7B). We observed that these NS4As expressed higher than ZIKV NS4A, and all were immunoprecipitated by ANKLE2-3xFLAG (Figure 8A). To further investigate this, we used Jensen-Shannon divergence to measure conservation at each amino acid position along NS4A (Figure 8B) (62). This showed regions of ANKLE2-interacting region (TM2 and TM3), including the C-terminus have high levels of conservation (Jensen-Shannon divergence score > 0.7). Sequence logo analysis of this region using a broader set of flaviviruses (all DENV serotypes, St. Louis encephalitis virus [SLEV], Powassan virus [POWV] and Langat virus [LGTV]) (63, 64) revealed several amino acids in the interaction domain that were highly conserved and could mediate the interaction with ANKLE2 (Figure 8C).

**Figure 8:**
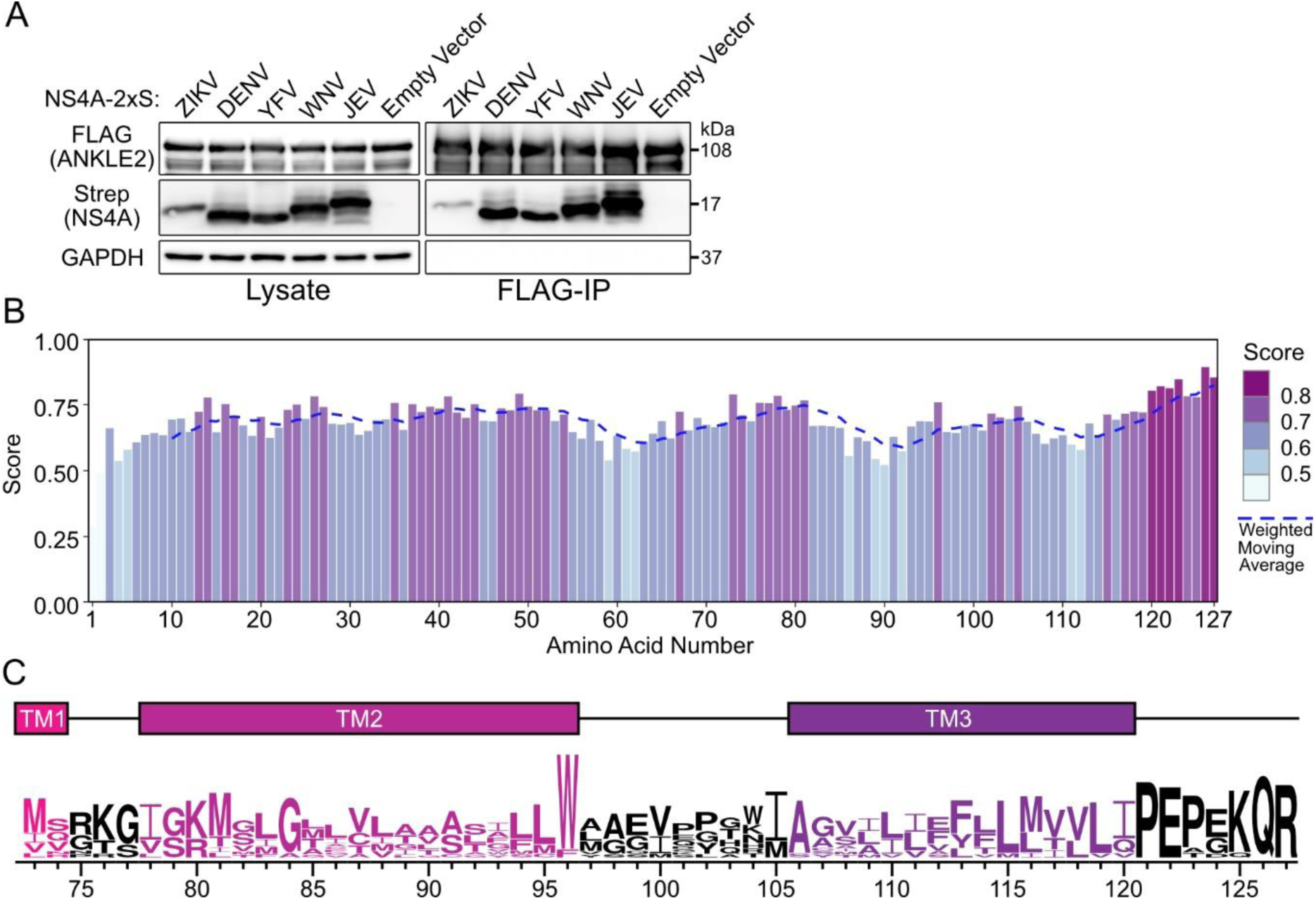
Conservation of flavivirus NS4A and its interaction with ANKLE2. **A)** Western blot of lysate and FLAG IP samples for indicated flavivirus NS4A protein. ANKLE2-3xFLAG co-transfected with an empty Strep plasmid was used as a control. **B)** Conservation of NS4A measured by Jensen-Shannon divergence at each amino acid position (62). **C)** Logo analysis of the ANKLE2 interaction domain of NS4A (TM2 and TM3, amino acids 73-127), compiled from 12 flaviviruses, with consensus TM domains from ZIKV NS4A overlayed (63, 64). Only a portion of TM1 is shown as a reference.

## Discussion

In this study we explore the relationship between ZIKV and host ANKLE2, with a focus on virus replication. Knockdown of ANKLE2 led to reductions in ZIKV replication at multiple timepoints across a range of MOIs and was present for multiple ZIKV strains. The observed decreases in ZIKV replication are modest (maximum effect ∼10-fold), likely due to the incomplete nature of our knockdown. It is possible that full genetic knockout of ANKLE2 would yield more dramatic virus replication defects.

The supportive role of ANKLE2 in ZIKV replication raises exciting possibilities about ANKLE2 function at sites of replication. Beyond BANF1, PP2A, and VRK1 (43, 44), ANKLE2 also interacts with many other host proteins. For example, ANKLE2 also influences the cell cycle by interacting with Aurora-A and ERα to mediate ERα phosphorylation (65). Plentiful other ANKLE2-host protein interactions have been found in large proteomic screens (66–69). Presumably the bulk of these interactions are mediated through its ANK domain, which facilitates scaffolding functions in many proteins (70, 71), or by the LEM domain, which is known to mediate interaction with the inner nuclear membrane lamina and BANF1 (72–74). It is particularly appealing to speculate that ZIKV leverages the protein interaction/scaffolding function of ANKLE2 to facilitate protein interactions within replication compartments. Future mechanistic studies exploring ANKLE2 protein interactions in replication compartments will be valuable.

To our knowledge, the NS4A-ANKLE2 interaction is a rare example of a ZIKV-host protein interaction that dysregulates the host function to cause neuropathogenesis (38, 40) in the process of promoting ZIKV replication. Several other protein interactions impact aspects of neurodevelopment but appear distinct from virus replication. ZIKV NS3 cleaves host BMP2, inducing osteogenesis and intracranial calcification commonly seen in CZS (27). Expression of NS2A *in vivo* impacts neurodevelopment by disrupting adherens junctions in radial glial cells (31). Expression of NS4A and NS4B impairs the growth of neural stem cells *in vitro* and perturbs autophagy (29). Yet the host factors involved in pathogenesis were not linked to virus replication directly in these studies. A notable exception is the interaction between ZIKV Capsid and Dicer. Capsid interacts with Dicer and inhibits its antiviral activity to promote ZIKV replication while simultaneously inducing neurodevelopmental defects (35). These types of virus-host interactions that result in compounding losses for the host represent an exciting model system to simultaneously study virus replication and pathogenesis.

Our work provides molecular-level insight into the biophysical nature of the NS4A-ANKLE2 interaction. Co-IP studies revealed that both the TM and LEM domains of ANKLE2 together mediate the interaction. Interestingly, loss of the TM alone is sufficient to disrupt the concentration of ANKLE2 to sites of NS4A during infection, though the biochemical interaction is still possible following deletion of just the TM domain. The TM anchors ANKLE2 to the ER and nuclear envelope, and its loss reduces this localization in favor of cytoplasmic localization (59). This deletion separates ANKLE2 from NS4A, which would otherwise interact with the LEM domain. This does not imply an inherent necessity of the TM domain for the interaction with NS4A. The presence of this domain varies between organisms with no clear phylogenetic separation. Mammalian (human, primate, mouse, etc.) ANKLE2 and the *C. elegans* homolog LEM-4L all contain a N-terminal TM. *Drosophila* Ankle2 lacks a clear TM but does have structured elements at the N-terminus (Supplementary Figure 5). Interestingly, *Drosophila* Ankle2 still localizes to the ER and is inhibited by NS4A (40), suggesting this structured region may serve as a previously uncharacterized TM, or it has another mechanism of ER localization. Moreover, the interaction with NS4A could still be detected biochemically by co-IP for the TM-deleted ANKLE2. This is likely due to mixing of cytoplasmic and ER compartments during lysis, which enables the interaction through the LEM domain even in the absence of correct ER localization. Thus, ER localization and the TM and LEM domains are the major contributors to the interaction. Dissecting this interaction with amino acid resolution in the future has the potential to identify ANKLE2 mutants that do not interact with NS4A (or vice versa). This would enable direct testing of the importance of this protein interaction for ZIKV replication. Ultimately it could reveal protective ANKLE2 variants which are functional in brain development, but do not interact with NS4A and thus do not support replication. Thus, this compounded loss scenario for the host could be turned into a compounded win

Finally, we identified the NS4A determinants of the interactions and demonstrate that the NS4A-ANKLE2 protein interaction is highly conserved amongst mosquito-borne flaviviruses. The conserved nature of the interactions suggests that while inhibition of ANKLE2 to cause microcephaly may be unique to ZIKV, coopting ANKLE2 to promote replication maybe a general feature of flaviviruses. By analyzing the sequence of this interaction domain (TM2 and TM3) for diverse flaviviruses, we were able to identify highly conserved amino acids in this region that may mediate the interaction with ANKLE2. Mutagenesis of this region will be critical to generating a ZIKV mutant that does not interact with ANKLE2. Such a mutant could be evaluated for replication, and if viable, neuropathogenesis.

In summary, we report that knockdown of *ANKLE2* reduces ZIKV replication. During infection, ANKLE2 becomes concentrated at sites of virus replication within the ER. The NS4A-ANKLE2 physical interaction is mediated by the N-terminal TM and LEM domains of ANKLE2, whereas NS4A requires TM2 and TM3. Future studies exploring how ANKLE2 function is hijacked by NS4A in virus replication compartments and which amino acids mediate this interaction will provide molecular mechanistic insight into this virus-host protein interaction involved in replication and pathogenesis.

## Methods

### Cells

HEK293T (gift of Dr. Sam Díaz-Muñoz), Huh7 (gift of Dr. Raul Andino), and Vero (ATCC) cell lines were maintained in Dulbecco’s modified Eagle’s medium (DMEM, Gibco ThermoFisher) supplemented with 10 % fetal bovine serum (FBS, Gibco ThermoFisher) at 37°C, 5% CO_2_. Cells we washed with Dulbecco’s phosphate buffered saline (D-PBS, Life Technologies) and dissociated with 0.05% Trypsin-EDTA (Life Technologies). Cells were tested for *Mycoplasma spp.* monthly by PCR.

### Plasmids

Lentiviral plasmid (pHR-UCOE-EF1α-KRAB-dCas9-P2A-Bls) encoding a catalytically dead Cas9 (dCas9) with a C-terminal hemagglutinin (HA) tag was a gift from Dr. Sean Collins. For stable, inducible expression in HEK293T cells, ANKLE2 and GFP sequences were amplified by PCR and cloned into pLVX-TetOne-Puro, cut with EcoRI, using Gibson assembly. Codon-optimized ANKLE1 sequence was acquired from Twist BioSciences and similarly inserted into pLVX-TetOne-Puro, cut with EcoRI, with C-terminal APEX2 and 3xFLAG affinity tags. ANKLE2 truncations were designed based on AlphaFold2 structural prediction (57, 58) using NM_015114.3 accession sequence (searched August 8^th^, 2021). Codon optimized DNA fragments with C-terminal 3xFLAG affinity-tags were acquired from Twist Bioscience and inserted into pcDNA4_TO, cut with KpnI and ApaI, using Gibson assembly. A full length pcDNA4_TO ZIKV NS4A-2xStrep plasmid was previously generated (38), and used as a template for generation of NS4A truncation sequences using PCR amplification. Products were inserted into pcDNA4_TO cut with BamHI and EcoRI upstream of 2xStrep using Gibson assembly. Flavivirus (DENV, YFV, WNV, JEV) NS4A sequences were acquired from Twist Bioscience and inserted into pcDNA4_TO, cut with BamHI and XhoI, with C-terminal 2xStrep tags using Gibson assembly. All plasmids were prepared in Stbl3 or DH5α using MiniPrep (Sigma-Aldrich) or MidiPrep (Macherey-Nagel) kits and verified using sequencing services provided by GeneWiz. All sequence accession numbers are available in Table S2. Primer sequences used for the generation of all our constructs are available in Table S3.

### Lentiviral packaging, transduction, and cell selection

Lentiviral packaging and transduction were performed as previously described (75) using the calcium phosphate protocol (76). In short, 3.5 µg of cloning product plasmid were transfected into HEK293T with lentiviral packaging plasmids including 1.8 μg pMDLg/p-RRE, 1.25 μg pCMV-VSV-g and 1.5 μg pRSV-Rev. After 48 hours lentivirus particles were collected, and cell debris was removed by centrifugation (Eppendorf centrifuge 5810 R, Rotor S-4-104, 94 g, 5 minutes) and filtration through a 0.45 μm filter. The resulting lentiviral stocks were used to transduce HEK293T cells. Transduced cells were bulk selected for puromycin resistance (10 μg/ml, ThermoFisher). A control lentiviral plasmid encoding GFP without a selection marker was used in tandem as a control to ensure efficient packaging, transduction, and selection.

### Viruses and stock preparation

All ZIKV stocks were propagated in Vero cells and monitored for CPE. Supernatant was then harvested, and cell debris was removed by centrifugation (Eppendorf centrifuge 5810 R, Rotor S-4-104, 211 g, 5 minutes, 4°C) Cleared supernatant was then distributed into 500 µL aliquots and frozen at -80°C. Each aliquot was only used once to prevent repetitive freeze-thaw. Aliquots were titered by plaque assay (method below). Strains used were ZIKV PLCal/2013 (gift of Dr. Richard Wozniak), ZIKV PRVABC59 (gift of Dr. Lark Coffey), and ZIKV MR766 (BEI Resources, NIAID, NIH, as part of the WRCEVA program: Zika Virus, MR 766, NR-50065).

### *ANKLE2* CRISPRi knockdown and ZIKV Infection

Custom synthetic guide RNAs (gRNA) were acquired from Sigma and resuspended to 3 µM in TE buffer (10 mM Tris Base, 1 mM EDTA, pH 8.0). For Huh7-dCas9 knockdown in 12-well dishes 30 µL of each gRNA was combined with 7 µL TransIT-CRISPR transfection reagent (Sigma) in 363 µL OPTI-MEM (Life Technologies) and complexed at room-temperature for 20 minutes prior to being added to each well (90nM final gRNA concentration). A total of 1.2×10^5^ Huh7-dCas9 cells were then added and grown overnight at 37°C. Additional DMEM was then added 24 hpt. Viability experiments were done 72 hpt with ZombieGreen dye (BioLegend, gift of Dr. Scott Dawson) diluted 1:100 in D-PBS and incubated on live cells for 5 minutes. Ten images were taken for each condition and total and dead cells were then counted to determine viability. ZIKV replication after *ANKLE2* knockdown was done by removing media from each well 72 hpt. 2 mL of fresh DMEM and appropriate volume of ZIKV stock was then added to each well. Aliquots of supernatant were harvested at 0, 18, 24, 48, and 72 hpi and frozen at –80°C.

### Plaque assay

Vero cells were grown as a monolayer in 6-well dishes overnight. Virus aliquots were thawed on ice then subjected to 10-fold serial dilution. Media was removed from Vero cells and the monolayer was washed once with 2mL D-PBS. 500µL of each virus dilution was then added and incubated for 1 hour at 37°C with periodic rocking. Virus was then removed and cells were overlayed with 3 mL of DMEM with 0.8% methylcellulose (Sigma), 1% FBS, 1% penicillin/streptomycin (Thomas Scientific) and incubated at 37°C for 4 days. Cells were then fixed with 4% formaldehyde (Fisher) for 30 minutes at room temperature. Formaldehyde and media were then removed, and cells were stained with 0.23% crystal violet solution (Fisher) for 30 minutes. Solution was then removed, and plaques were counted.

### Quantitative RT-PCR

RNA was harvested using *Quick*-RNA Miniprep kit per manufacturer’s instructions (Zymo). Purified RNA (500ng) was then used to make cDNA using iScript^TM^ cDNA synthesis kit (Bio-Rad). After cDNA synthesis each sample was resuspended to a total volume of 100µL using RNase-Free water. A total of 2 µL of cDNA was then used for each qPCR reaction (done in technical triplicates) using LightCycler 480 SYBR Master Mix (Fisher). Samples were run in a Roche LightCycler® 480 II Instrument. Changes in gene expression were calculated using the Livak method (2^ΔΔCt^) (77).

### Confocal immunofluorescence microscopy

HEK293T or Huh7 cells cultured on #1.5 coverslips were fixed with 4% paraformaldehyde (Fisher) for 15 minutes at room temperature. Cells were permeabilized with 0.1% Triton X-100 (Integra) for 10 minutes and blocked with 5% goat serum (Sigma) in PBS-Tween (0.1% Tween-20, Fisher). Coverslips were incubated with primary antibodies overnight at 4°C. Coverslips were then washed in PBS-Tween and incubated in secondary antibody at room temperature for 1 hour. Nuclei were visualized with Hoechst (Invitrogen). Confocal images were acquired using an Olympus FV1000 Spectral Scan point-scanning confocal fitted to an Olympus IX-81 inverted microscope using a PlanApo 60x/NA1.40 oil immersion lens or Zeiss Airyscan LSM800 with Axiocam using a 63x/NA1.40 oil immersion lens. Laser lines at 405, 488, and 543nm were employed sequentially for each image using optics and detector stock settings in the “Dye List” portion of the FluoView microscope-controlling software. All antibodies and dilutions are listed in Supplementary Table S1. Confocal images were analyzed using ImageJ (Fiji) software (78). Signal colocalization was quantified using Pearson’s correlation coefficient (R-value) determined with the “Colocalization 2” analysis tool within Fiji after masking individual cells across at least 5 images.

### ANKLE2 and NS4A co-transfection and FLAG affinity-purification

For transfection 5×10^6^ HEK293T cells were plated in 10 cm dishes and grown overnight. Transfection was performed by combining 3.5 µg of each corresponding plasmid DNA with 700 µL of serum-free DMEM. Next, 21 µL of PolyJet transfection reagent (SignaGen) was combined with 700 µL serum-free DMEM and added to each plasmid DNA tube. Samples were mixed and incubated at room temperature for 15 minutes prior to addition to cells. Cells were then grown for an additional 24 hours. Transfection efficiency was confirmed using a GFP encoding plasmid. Media was then removed from each plate. To dissociate cells, 5 mL of D-PBS supplemented with 10 mM EDTA was added and allowed to incubate for several minutes. Cells were resuspended in 5 mL of D-PBS and transferred to 15 mL conical tubes prior to centrifugation 94 g, 4°C for 5 minutes (Eppendorf centrifuge 5810 R, Rotor S-4-104). Cell pellets were washed with 5 mL D-PBS and centrifugation was repeated. Supernatant was removed and pellets were then resuspended in 1 mL IP buffer (50 mM Tris Base, 150 mM NaCl, 0.5 M EDTA, pH 7.4) with Pierce^TM^ protease (Thermo Scientific) and Pierce^TM^ phosphatase (Fisher) inhibitor tablets supplemented with 0.5% NP-40 Substitute (Igepal^TM^ CA-630, Affymetrix). Cells were lysed for 30 minutes at 4°C, and lysate was then centrifugated at 845 g, 4°C, for 20 minutes (Eppendorf centrifuge 5424 R, Rotor FA-45-24-11). A portion of each lysate (60-100 µL) was collected, normalized by BCA assay (Thermo Scientific), and saved for western blot analysis. Remaining lysate was added to 40 µL of magnetic FLAG beads (Sigma) and incubated overnight at 4°C with gentle rotation. Beads were then washed 4 times with 1 mL IP buffer with 0.05% NP-40 and once with 1 mL IP buffer without NP-40. Beads were then incubated in 40 µL of 100 ng/mL 3x FLAG peptide (APExBIO) at 211 g for 1 hour at room temperature (Eppendorf ThermoMixerC). Eluate was then removed. Eluate and lysate were resuspended in NuPAGE LDS sample buffer and bond-breaker TCEP (Thermo Scientific) according to manufacturer’s recommendation. Samples were boiled for 10 minutes at 95°C prior to evaluation by western blot (below).

### Subcellular fractionation

Proteins were isolated from the cytosol, membrane-bound organelles, and the nucleus using a previously established protocol for cultured cells (79). In brief, cells were sequentially lysed in buffer (3 M NaCl, 1 M HEPES, 1 M glycerol, 1X protease inhibitor) containing increasingly stronger detergents. Following cell trypsinization, buffer with digitonin (25 µg/mL) disrupted the plasma membrane over the course of gentle rotation at 4°C. Buffer with Igepal (1% v:v, NP-40 substitute) permeabilized membrane-bound organelles during incubation on ice. The nuclear membrane was disrupted by buffer containing sodium deoxycholate (0.5% w:v) and sodium dodecyl sulfate (0.1 % w:v), with subsequent sonification to disrupt genomic DNA. All separations were performed by centrifugation (Eppendorf centrifuge 5424 R, Rotor FA-45-24-11, 4°C). Cell fractions were evaluated by western blot.

### Western blot

For whole cell lysates, cells were lysed in RIPA buffer (150 mM NaCl, 50 mM Tris Base, 1% Triton X-100, 0.5% sodium deoxycholate) supplemented with protease inhibitors for 5 minutes at room temperature. Cell lysate was incubated on ice for 30 minutes prior to centrifugation (Eppendorf centrifuge 5424 R, Rotor FA-45-24-11, 13,500 g, 4°C, 20 min). When possible, the total protein concentration of each sample was normalized by BCA assay. Protein samples (lysates or IP eluates) were resuspended in NuPAGE LDS sample buffer supplemented with TCEP and boiled at 95°C for 10 minutes. Samples were run on 7.5-12% polyacrylamide gels for ∼1 hour at 150V and transferred to PVDF membranes (VWR) for 1 hour at 330 mA on ice. Membranes were then blocked in 5% milk solution for 1 hour prior to overnight incubation in primary antibodies (Table S1) at 4°C. Membranes were washed 3x in Tris-buffered saline with Tween-20 (TBS-T) (150 mM NaCl, 20 mM Tris Base, 0.1% Tween-20, Fisher) and incubated with HRP-conjugated secondary antibodies in 5% milk for 1 hour at room temperature. Membranes were again washed 3x in TBS-T and 1x in TBS (without Tween-20) prior to Pierce^TM^ ECL activation (Fisher). Membranes were imaged using Amersham Imager 600 (GE). Western blot images were analyzed using Fiji. Densitometry was calculated by measuring the band intensity ratio of the experimental band to the loading control band.

### Statistical analysis

Statistical analysis and plotting were performed using GraphPad Prism 6 software (GraphPad Prism 6.0; GraphPad Software Inc., La Jolla, CA, USA). Error bars represent standard deviations. Data were considered statistically significant when a p < 0.05 was determined by Student’s T-test or one-way ANOVA with noted multiple-comparison test.

## Acknowledgements

We would like to thank members of the Shah lab for their constant support and helpful feedback. Funding was provided by the University of California, Davis and the W. M. Keck foundation. ATF was partially supported by NIH T32 fellowships (5T32GM7377-42 and 2T32AI060555-16). STH was partially supported by NIH T34 (T34GM136469). The Olympus FV1000 confocal used in this study was purchased using NIH Shared Instrumentation Grant 1S10RR019266-01. We thank the MCB Light Microscopy Imaging Facility, which is a UC Davis Campus Core Research Facility, for the use of this microscope. We thank Dr. Neil Hunter’s lab for the use of their AiryScan Confocal microscope.

## Author Contributions

ATF and PSS conceived of work and designed experiments. ATF performed and analyzed all experiments. MWK, VH, NL, TNS, and STH assisted with experiments. STH, NL, TNS, and SK performed cloning and plasmid preparation. ATF and PSS wrote the manuscript. All authors revised the manuscript. PSS secured funding for work. All authors read and approved the final manuscript.

---

**Supplementary Figure 1:**
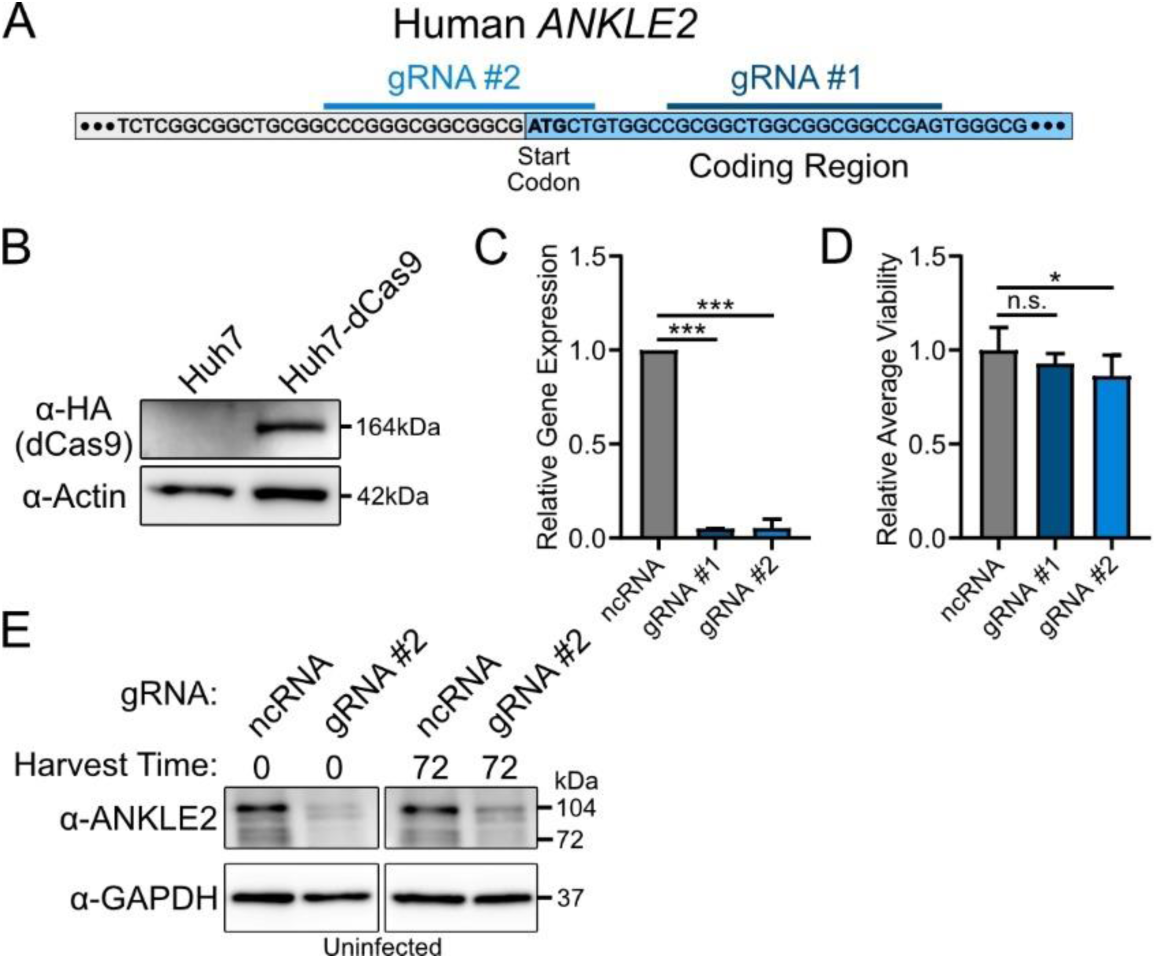
Knockdown of *ANKLE2* in Huh7-dCas9 cells using CRISPRi. **A)** Schematic of *ANKLE2* and the targeting guide RNAs (gRNA) that were used to knockdown *ANKLE2* expression. **B)** Western blot to evaluate the expression of dCas9-HA in Huh7 cells that were used for CRISPRi experiments. **C)** Western blot to evaluate ANKLE2 expression after CRIPSRi knockdown with three different gRNAs. The primary isoform is shown at 104kDa, and the knockdown of various other isoforms can be observed at 76 and 72kdA. **D)** Densitometry analysis to quantify ANKLE2 band intensity. ANKLE2 band intensity was normalized to its respective loading control and then normalized to ncRNA. Error bars represent the standard deviation between 3 biological replicates. **E)** *ANKLE2* gene expression after knockdown was quantified by qRT-PCR. Error bars represent the standard deviation between 3 biological replicates. **F)** Cell viability was evaluated using ZombieGreen dye. Data represent the average cell viability across 10 images for each condition. **D-F)** Error bars represent the standard deviation. Student’s unpaired two-tailed t-test, compared to ncRNA, n.s., not significant, * p > 0.05, *** p > 0.001.

**Supplementary Figure 2:**
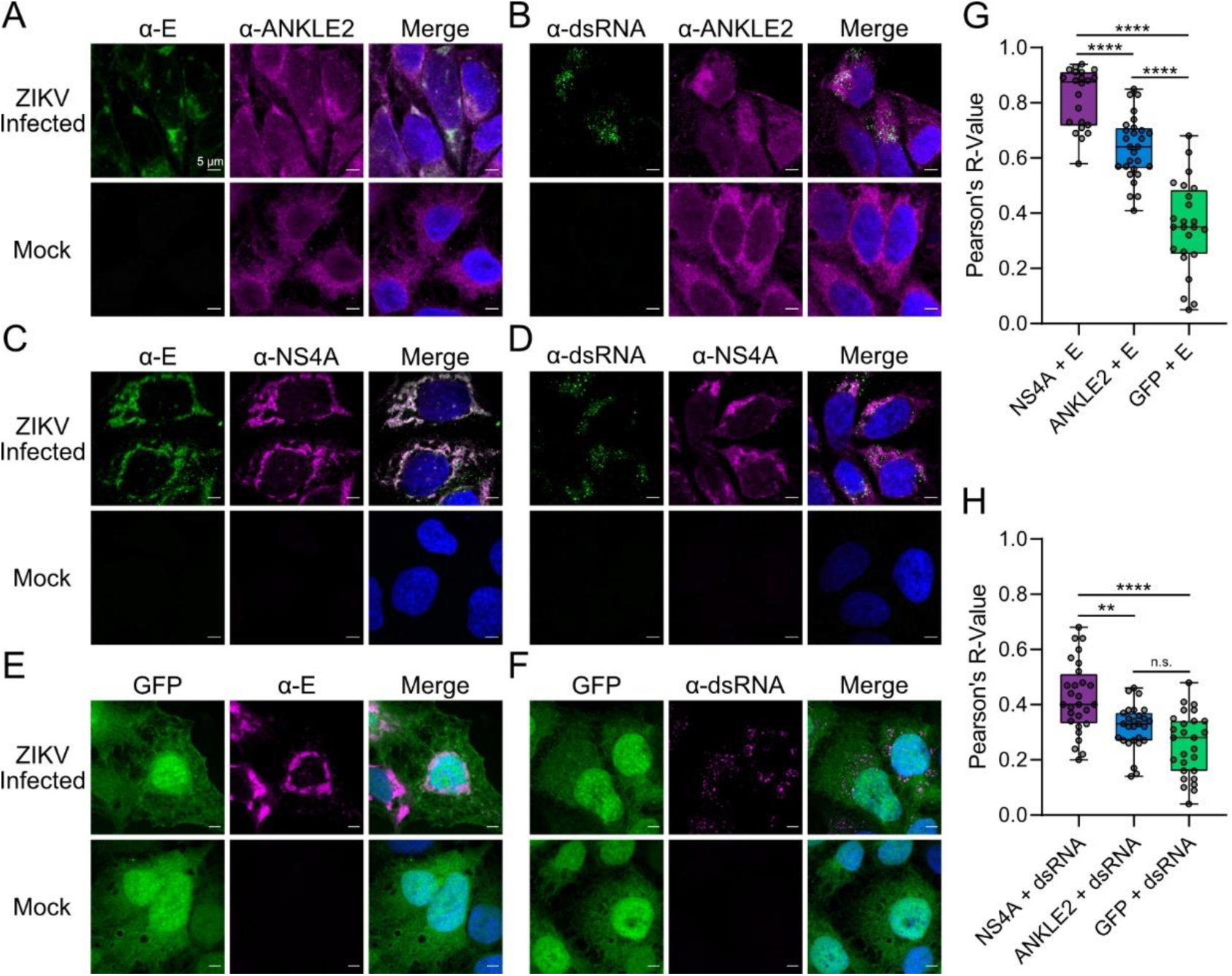
ANKLE2 colocalizes with sites of ZIKV genome replication in Huh7-dCas9 cells. **A-F)** Huh7-dCas9 cells plated on cover slips were infected with ZIKV for 24-48 hours and subsequently immunostained for with noted antibodies and their target (in the case of 4G2/Env and rJ2/dsRNA). Cover slips were imaged by confocal microscopy and analyzed using ImageJ. All scale bars = 5 µm **G-H)** Pearson’s R-value measured using ImageJ colocalization tool. N = 22 to 28 cells across 5 images per condition. One-way ANOVA with Tukey’s multiple comparisons test, n.s. = not significant, ** p < 0.01, **** p < 0.0001.

**Supplementary Figure 3:**
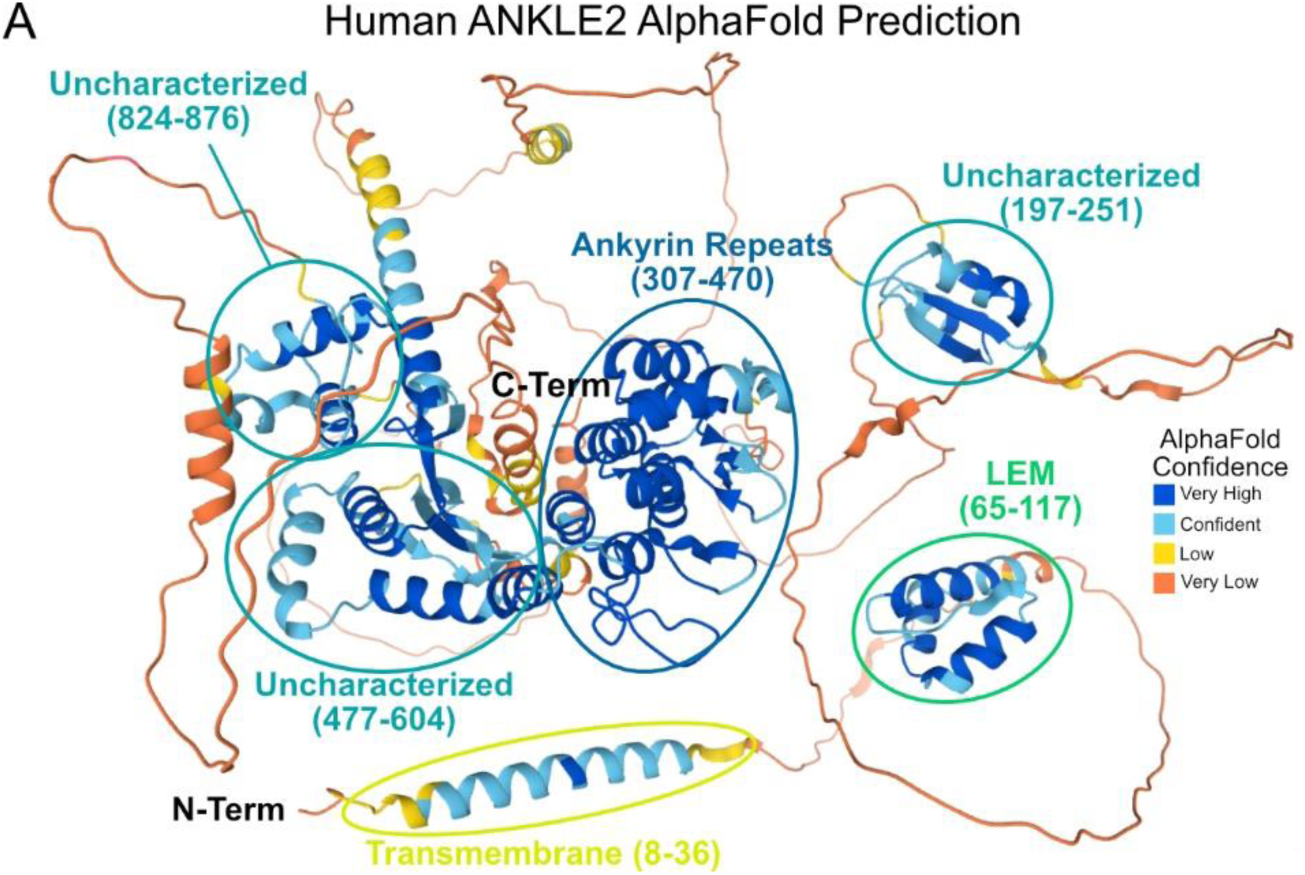
AlphaFold2 protein structure prediction reveals uncharacterized structural regions in human ANKLE2. AlphaFold2 was searched for human ANKLE2 and predicted structures were compared against known domains. AlphaFold2 confidence is shown as a gradient from very high (blue) to very low (orange). Known domains are highlighted, ankyrin repeats = dark blue, LEM = green, TM = yellow, uncharacterized = light blue.

**Supplementary Figure 4:**
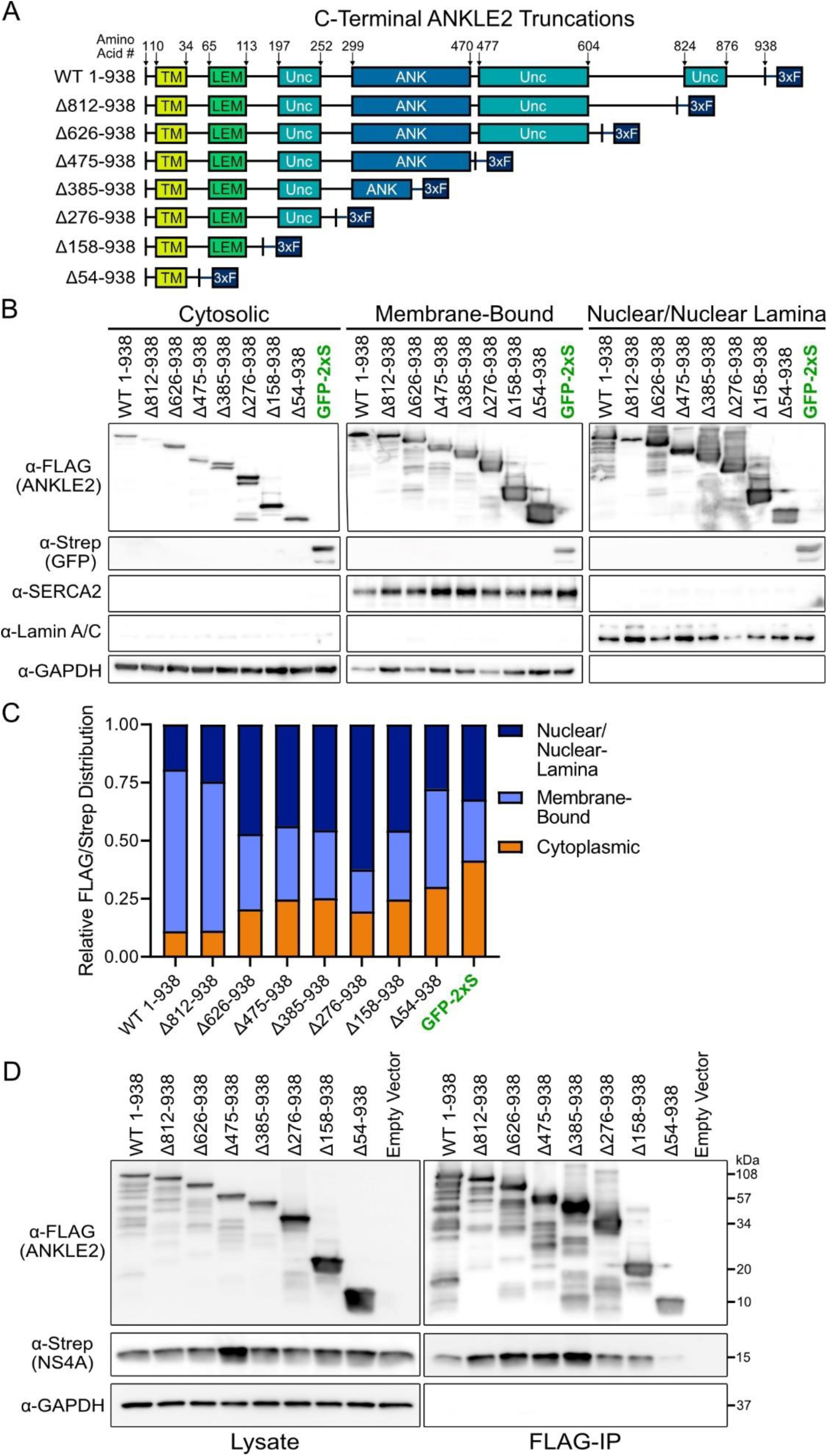
ANKLE2 C-terminal truncations interact with ZIKV NS4A and are retained primarily in the ER and Nuclear regions. **A**) C-terminal ANKLE2 truncations were designed based on a structural prediction by AlphaFold2. TM = transmembrane domain; LEM = LAP2, emerin, MAN1 domain; Unc = uncharacterized structure predicted by AlphaFold2; ANK = ankyrin repeat domain; 3xF = 3x FLAG affinity tag. **B**) Western blot analysis of subcellular fractions of cells transfected with the indicated ANKLE2 construct. SERCA2 and Lamin A/C antibodies were used to validate enrichment of membrane-bound and nuclear fractions, respectively, while GAPDH was used as a cytoplasmic marker. **C**) Densitometry analysis was performed on western blot images by measuring the intensity of FLAG/Strep bands relative to each fraction’s marker. These values were then plotted as a fractional distribution for relative intensity across all fractions. D) Western blot of lysate and FLAG-IP samples for indicated ANKLE2 construct cotransfected with NS4A.

**Supplementary Figure 5:**
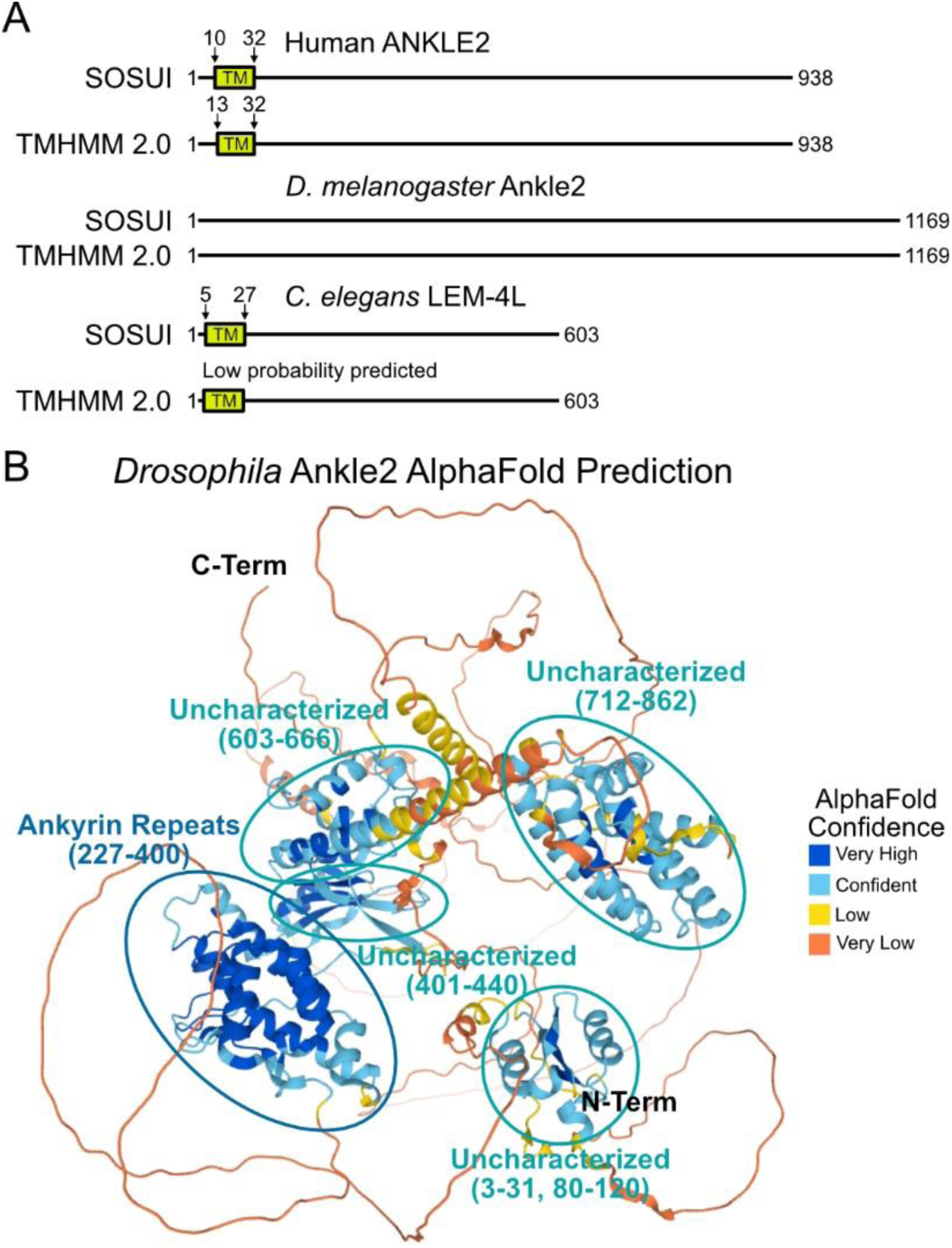
Transmembrane domain prediction of ANKLE2 homologs and structural prediction of *Drosophila* Ankle2. **A)** Transmembrane domain prediction of human ANKLE2, *Drosophila melanogaster* Ankle2 (NM_001298446.1), and *C. elegans* LEM-4L. TM prediction was determined using SOSUI and TMHMM 2.0 modeling (80, 81). **B)** AlphaFold2 was searched for *Drosophila* Ankle2 and predicted structures were compared against known domains. AlphaFold2 confidence is shown as a gradient from very high (blue) to very low (orange). Known domains are highlighted, ankyrin repeats = dark blue, uncharacterized = light blue.

**Supplementary Table 1:** Antibodies

| Antibody | Host Species | Dilution Used | Supplier (Catalog #) | RRID |
| --- | --- | --- | --- | --- |
| GAPDH | Mouse | 1:1000 (WB) | Fisher (PIMA515738) | AB_2537652 |
| FLAG-M2 | Mouse | 1:200 (IF),<br>1:1000 (WB) | MilliporeSigma (F1804) | AB_262044 |
| Strep | Mouse | 1:1000 (IF, WB) | Qiagen (34850) | AB_2810987 |
| 4G2 (Flavivirus E) | Mouse | 1:250 (IF) | ATCC (HB-112) | CVCL_J890 |
| rJ2 (dsRNA) | Mouse | 1:40 (IF) | MilliporeSigma (MABE1134) | AB_2819101 |
| Lamin A/C (4C11) | Mouse | 1:1000 (WB) | Cell Signaling (4777)<br>Gift from Dr. Jodi Nunnari | AB_10545756 |
| SERCA2 | Mouse | 1:100 (IF),<br>1:500 (WB) | Invitrogen (MA3919) | AB_325502 |
| ANKLE2 | Rabbit | 1:80 (IF) | Atlas Antibodies (HPA074838) | N/A |
| ANKLE2 | Rabbit | 1:1000 (WB) | Atlas Antibodies (HPA003602) | AB_1858349 |
| ZIKV NS4A | Rabbit | 1:1000 (IF, WB) | GeneTex (GTX133704) | AB_2887067 |
| Actin | Rabbit | 1:5000 (WB) | Sigma (A2066) | AB_476693 |
| Anti-Mouse IgG-HRP | Rabbit | 1:5000 (WB) | SouthernBiotech (6170-05) | AB_2796243 |
| Calreticulin | Rabbit | 1:500 (IF) | Abcam (AB92516) | AB_10562796 |
| Calnexin | Rabbit | 1:250 (IF) | Proteintech (10427-2-AP)<br>Gift from Dr. Jodi Nunnari | AB_2069033 |
| HA | Rat | 1:2000 (WB) | MilliporeSigma (3F10)<br>Gift from Dr. Joanna Chiu | AB_2314622 |
| Anti-Rabbit IgG-HRP | Goat | 1:5000 (WB) | SouthernBiotech (4030-05) | AB_2687483 |
| Anti-Mouse AlexaFlour-488 | Goat | 1:1000 (IF) | Invitrogen (A28175) | AB_2536161 |
| Anti-Rabbit AlexaFlour-555 | Goat | 1:1000 (IF) | Invitrogen (A27039) | AB_2536100 |
| Anti-Mouse AlexaFlour-680 | Goat | 1:1000 (IF) | Invitrogen (A21057) | AB_2535723 |

WB = Western blot, IF = Immunofluorescence

**Supplementary Table 2:** Sequences

| Sequence | Accession Number | Relevant Figure(s) |
| --- | --- | --- |
| Human ANKLE2 | NM_015114.3 | Figures 1, 2, 4, 5, 7 and 8<br>Supplementary Figures 1, 3, 4, and 5 |
| *D. melanogaster* Ankle2 | NM_001298446.1 | Supplementary Figure 5 |
| *C. elegans* LEM-4L | NM_001028343.3 | Supplementary Figure 5 |
| Human ANKLE1 | NM_152363.6 | Figure 2 |
| eGFP | UDY80669.1 | Figures 2, 3, 4, and 6<br>Supplementary Figures 2, and 4 |
| Zika virus (ZIKV) | MK713748.1 | Figures 5, 6, 7, and 8.<br>Supplementary Figure 4 |
| Dengue virus serotype 1 (DENV-1) | QZN14214.1 | Figure 8 |
| Dengue virus serotype 2 (DENV-2) | NC_001474.2 | Figure 8 |
| Dengue virus serotype 3 (DENV-3) | QXM02604.1 | Figure 8 |
| Dengue virus serotype 4 (DENV-4) | QTX92397.1 | Figure 8 |
| Yellow Fever virus (YFV) | KF769016.1 | Figure 8 |
| West Nile virus (WNV) | DQ211652.1 | Figure 8 |
| Japanese encephalitis virus (JEV) | NC_001437.1 | Figure 8 |
| Saint Louis encephalitis virus (SLEV) | QVN47903.1 | Figure 8 |
| Tick-borne encephalitis virus (TBEV) | JX498940.1 | Figure 8 |
| Powassan virus (POWV) | L06436.1 | Figure 8 |
| Langat virus (LGTV) | NC_003690.1 | Figure 8 |

**Supplementary Table 3:**
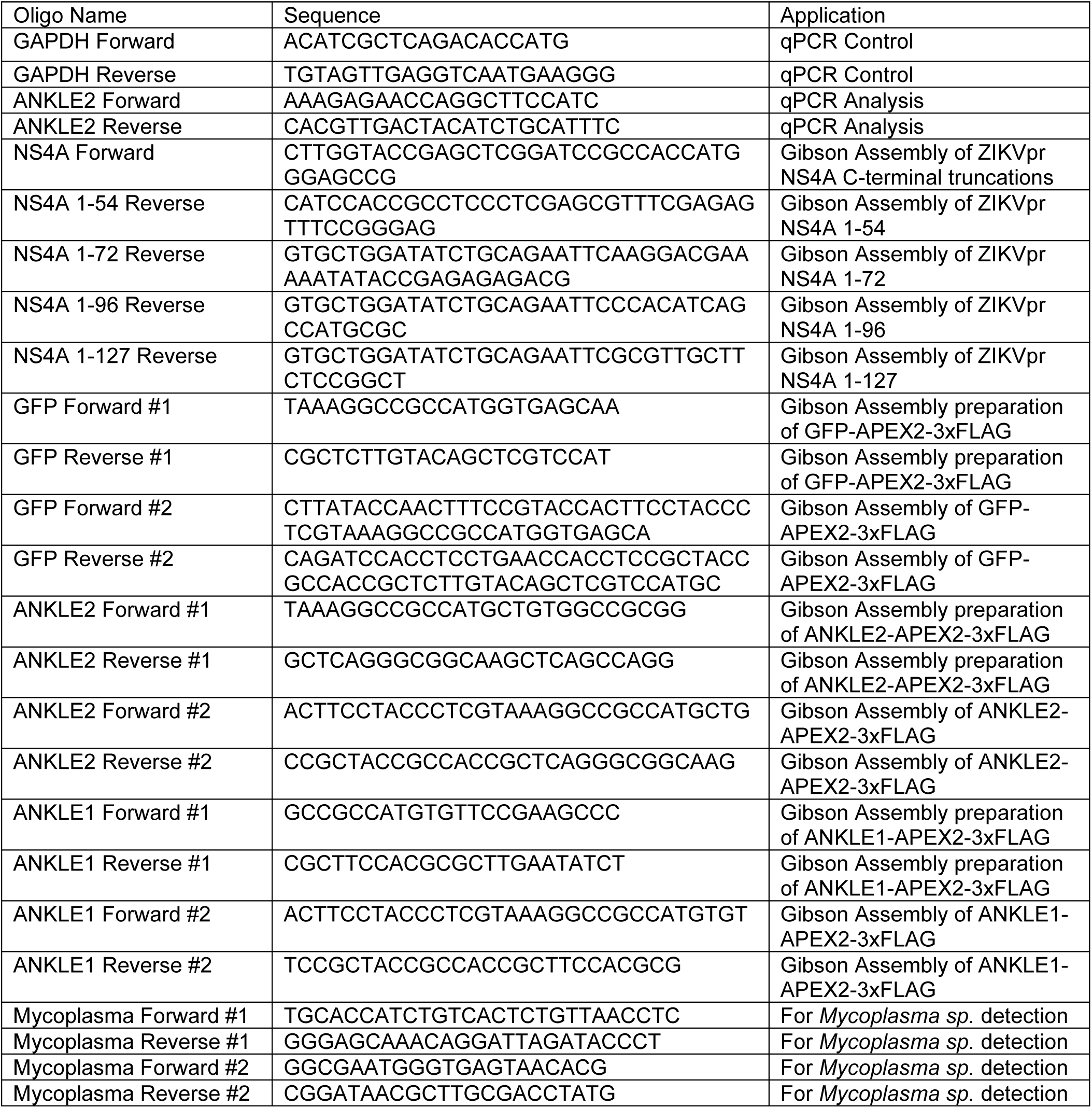
Oligonucleotides

